# Genetically Defined, Syngeneic Organoid Platform for Developing Combination Therapies for Ovarian Cancer

**DOI:** 10.1101/2020.04.06.028597

**Authors:** Shuang Zhang, Sonia Iyer, Hao Ran, Igor Dolgalev, Wei Wei, Connor Foster, Robert A. Weinberg, Benjamin G. Neel

## Abstract

The paucity of genetically informed, immune-competent tumor models impedes evaluation of conventional, targeted, and immune therapies. By engineering mouse fallopian tube (FT) organoids using lentiviral gene transduction and/or CRISPR/Cas9 mutagenesis, we generated multiple high grade serous ovarian carcinoma (HGSOC) models exhibiting mutational combinations seen in patients. Detailed analysis of homologous recombination (HR)-proficient (*Tp53*^*-/-*^;*Ccne1*^*OE*^;*Akt2*^*OE*^; *Kras*^*OE*^), HR-deficient (*Tp53*^*-/-*^;*Brca1*^*-/-*^;*Myc*^*OE*^) and unclassified (*Tp53*^*-/-*^;*Pten*^*-/-*^;*Nf1*^*-/-*^) organoids revealed differences in *in vitro* properties and tumorigenicity. Tumorigenic organoids had variable sensitivity to HGSOC chemotherapeutics and evoked distinct immune microenvironments. These findings enabled development of a chemotherapy/immunotherapy regimen that yielded durable, T-cell dependent responses in *Tp53*^*-/-*^;*Ccne1*^*OE*^;*Akt2*^*OE*^;*Kras* HGSOC; by contrast, *Tp53*^*-/-*^;*Pten*^*-/-*^;*Nf1*^*-/-*^ tumors failed to respond. Genotype-informed, syngeneic organoid models could provide an improved platform for rapid evaluation of tumor biology and therapeutics.

**HIGHLIGHTS:**

- Orthotopic injection of genetically defined fallopian tube organoids yield HGSOC.
- Ovarian tumors with different genotypes evoke distinct immune microenvironments
- Combining Gemcitabine, anti-PD-L1, and anti-CTLA-4 result in complete responses in *Tp53*^*-/-*^;*Ccne1*^*OE*^;*Akt2*^*OE*^;*Kras*^*OE*^ organoid-derived HGSOC
- Therapeutic response is tumor genotype-specific

## INTRODUCTION

The past 30 years of cancer research have yielded remarkable therapeutic advances along two main fronts (Esteva et al., 2019; Robert et al., 2016). “Targeted therapies” were developed against oncogenic “driver” tyrosine and serine/threonine kinases or key downstream signaling components (Otto and Sicinski, 2017). Concomitantly, powerful new “immune therapies” emerged, including cell therapies and “immune checkpoint inhibitors” (e.g., anti-CTLA4, anti-PD1, anti-PDL1), (Pardoll, 2012). These new modalities complement or replace conventional chemo- and radiation therapy, and are lifesaving for some patients. Nevertheless, most patients with metastatic solid tumors still succumb to their disease.

Targeted and immune therapies developed in parallel, usually use distinct experimental systems. Even today, targeted agents typically are tested against cancer cell lines/cell-derived xenografts (CDXs), patient-derived xenografts (PDXs), and/or more recently, human tumor spheroid/organoids. Advantages of these models include their human origin, relevant mutational/epigenetic events, and retention of some degree of tumor heterogeneity. However, such systems do not enable evaluation of anti-tumor immune responses. PDXs have been established in “humanized” mice, but ∼30% of human/mouse growth factors, cytokines, and chemokines fail to interact with the cognate receptor(s) in the other species, imposing an intrinsic limit on “humanization” (Walsh et al., 2017). Immune therapies, by contrast, are mainly tested against syngeneic mouse tumors (Mosely et al., 2017). These models (e.g., B16, CT26, MC38) are mainly carcinogen-induced, arise from unknown, irrelevant, or not the most relevant cell-of-origin, and often lack the key causal mutations found in the corresponding human disease. Some targeted agents/immune therapies have been evaluated in genetically engineered mouse models (GEMMs), which are designed to harbor disease-relevant genetic abnormalities and have intact immune systems (Kersten et al., 2017). Typically, only a few mutational combinations are generated for a given malignancy, limiting the diversity of the human disease that can be analyzed. Most GEMMs also introduce cancer-associated defects simultaneously into all epithelial cells in the target tissue. Real-world tumors, by contrast, initiate clonally and expand and progress in a sea of predominantly normal cells. A series of transplantable GEMM-derived melanoma models (Yum/Yummer) has been generated (Meeth et al., 2016), but these are based on the same truncal mutations with limited genetic diversity.

The tumor genotype, in the specific context of its cell-of-origin, determines susceptibility to conventional and targeted therapies, intrinsic immunogenicity (e.g., by neoantigens, altered surface expression of MHC class I molecules and/or ligands for activating/inhibitory receptors on immune cells), and the spectrum of cytokines and chemokines (“secretome”) that it produces (Binnewies et al., 2018; Li and Stanger, 2019; Wellenstein and de Visser, 2018). The secretome, in turn, initiates immune cell recruitment in the tumor microenvironment (TME). Save for mutation-selective agents (e.g., RAS^G12C^ inhibitors, osimertinib for mutant EGFR), targeted and conventional agents affect cells in the TME as well as tumor cells (Canon et al., 2019; Hallin et al., 2020). Therefore, a suite of immune-competent mouse models bearing tumors with frequently co-occurring genetically defects seen in patient neoplasms could enable more accurate evaluation of novel therapeutic agents or new combinations of existing drugs.

We chose to develop such a platform for high grade serous ovarian cancer (HGSOC). The most common and deadly form of ovarian epithelial cancer, causing ∼70% of deaths (Narod, 2016), HGSOC usually presents at an advanced stage. Typically, patients have bulky metastatic spread throughout the peritoneum, although some have more discrete tumor deposits. Current treatment is surgical “debulking” and platinum/taxane-based chemotherapy, and often results in complete responses (CRs). Nevertheless, disease almost always recurs, eventually in drug-resistant form. Despite the recent addition of anti-angiogenics (Avastin) and PARP inhibitors to the HGSOC armamentarium, survival has improved only marginally in the past 30 years, and most (70-90%) patients die from their disease (Bowtell et al., 2015). Clearly, better therapeutic strategies are needed for this deadly malignancy.

Much is known about the cellular and molecular pathogenesis of HGSOC. Despite its appellation, HGSOC most often initiates with mutation, deletion, or silencing of *TP53* in fallopian tube epithelium (FTE), not the ovary. The Cancer Genome Atlas (TCGA) reveals additional pathogenic single nucleotide variants, but HGSOC is primarily a disease of copy number abnormalities (CNAs), including amplifications, deletions, and more complex chromosomal rearrangements, which affect multiple genes and pathways (Cancer Genome Atlas Research, 2011). The most clinically useful molecular classification groups HGSOC cases by homologous recombination (HR) status. Alterations in known HR genes, including *BRCA1, BRCA2, RAD51*, or other Fanconi Anemia genes, occur in ∼40% of cases; another ∼15-20% have *PTEN* loss or *EMSY* amplification and are probably HR-deficient (Konstantinopoulos et al., 2015). Defective HR confers sensitivity to platinum chemotherapy (the mainstay of HGSOC treatment), and some (but not all) of these defects also confer PARP inhibitor responsiveness (Ashworth, 2008; Franzese et al., 2019). The remaining ∼40% of tumors are HR-proficient, respond poorly to current therapy, and result in shorter survival (Nakayama et al., 2010). *CCNE1* amplification, found in ∼20% of cases, is notorious for causing chemo-resistance and poor outcome (Au-Yeung et al., 2017); hence, there is a particular need to develop new therapeutic strategies for these tumors. Despite the impressive progress in delineating the molecular anatomy of HGSOC, how specific combinations of mutations determine the transformed phenotype, including the tumor transcriptome, host immune response, and therapy response, remains poorly understood.

The paucity of genetically relevant, immune-competent models of HGSOC poses a major barrier to new therapeutic development. Many studies have used conventional cancer cell lines, most of which (including the most frequently used) lack the characteristic genomic abnormalities seen in HGSOC (Domcke et al., 2013). Human HGSOC organoids have been derived, but, as noted above, they cannot model anti-tumor immune responses (Hill et al., 2018; Kopper et al., 2019). ID8 cells and their derivatives are the mainstay of immunological studies of HGSOC, but these cells originate from ovarian surface epithelium, not FTE, and have wild type (WT) *Tp53* (Cancer Genome Atlas Research, 2011; Roby et al., 2000). Some HGSOC GEMMs that use FTE-selective promoter/enhancers to direct mutational events have been developed, but these involve artificial alterations (e.g., SV40 large T antigen expression) or mutational combinations (e.g., *Brca1*/*Pten/Tp53*) found relatively rarely in HGSOC, and most are on mixed strain backgrounds, which impedes some tumor immunology studies. Notably, no immune competent models for *CCNE1*-amplified HGSOC have been reported.

Capitalizing on our recently developed conditions for culturing normal and neoplastic mouse FTE organoids (Zhang et al., 2019), combined with viral-based overexpression and CRISPR/Cas9 mutagenesis, we developed multiple new syngeneic models of HGSOC. We demonstrate the utility of this platform for uncovering cellular genotype/phenotype relationships, complementation groups for tumorigenicity, the effect of tumor genotype on drug sensitivity, secretome, tumor immune landscape, and pattern of metastatic spread and, most importantly, the rational development of a highly effective therapeutic combination for *Ccne1*-overexpressing HGSOC using existing combinations of FDA-approved drugs.

## RESULTS

### FTE organoid-based platform for modeling HGSOC

Most cases of HGSOC initiate from the distal fallopian tube (fimbria), which is composed mainly of secretory (PAX8+) and ciliated (acetyl-α-tubulin+) cells (Ducie et al., 2017; Labidi-Galy et al., 2017). The initial event (except in patients with germ line mutations of *BRCA1/2* or other predisposing genes) is mutation of *TP53* in a PAX8+ cell, which, in concert with other defects, results in a precursor lesion, termed serous tubal intraepithelial carcinoma (STIC). TCGA revealed multiple combinations of SNVs/CNAs in HGSOC; superimposed on mutant *TP53*, these additional mutational events confer invasive potential and promote metastasis to the ovarian surface and peritoneum (Perets and Drapkin, 2016; Rebbeck et al., 2002).

We used our FTE organoid culture system (Zhang et al., 2019) to model the complex biology of HGSOC. Briefly, fimbrial cells from *Tp53*^*f/f*^ (or, where indicated, *Brca1*^*f/f*^:*Tp53*^*f/f*^ mice; see below) were seeded into Matrigel and cultured in defined media. Cyst-like organoids formed from single PAX8+ cells, a mixture of secretory and ciliated (acetyl-α-tubulin+) cells was seen after 6 days of culture, and tube-like epithelial folds developed by 10 days (Zhang et al., 2019) (Figure S1, and data not shown). After expansion, floxed alleles were excised by infection with adenovirus-Cre (Ad-Cre), yielding parental *Tp53*^*-/-*^ organoids (Figure S2A) or, where indicated, compound mutant mice (all in C57BL6/J background). Additional genetic changes were introduced by lenti- or retroviral gene transduction (to model over-expression) and/or CRISPR/Cas9 mutagenesis (to model deletions or mutations). These models were tested in various cellular assays or transferred to 2D cultures for larger scale studies. Tumorigenesis was assessed by orthotopic injection into the ovarian bursa (for details, see Experimental Procedures). Our current collection of models and their tumorigenic potential are summarized in Table S1. To evaluate the utility of this platform for studying HGSOC pathogenesis and therapeutics, representative examples of HR-proficient, HR-deficient, and unclassified subgroups were studied in greater detail.

### *Tp53*^*-/-*^; *Brca1*^*-/-*^;*Myc*^*OE*^ FTE organoids give rise to HGSOC-like tumors

*BRCA1/2* gene alterations are found in ∼20% of HGSOC (Kuchenbaecker et al., 2015), so we generated *Tp53*^*-/-*^*/Brca1*^*-/-*^ models to represent the HR-deficient subgroup (Figure 1A and Table S1). We infected *Tp53* ^*f/f*^;*Brca1* ^*f/f*^ FTE with Ad-Cre, picked single organoids, and confirmed deletion of the relevant loci (Figure 1B and C). Neither *Tp53* nor *Brca1* deletion alone or in combination grossly altered organoid morphology or significantly affected ciliated cell differentiation (Figure 1D and E). *Tp53*^*-/-*^;*Brca1*^*-/-*^ organoids were significantly larger than their parental counterparts, most likely due to increased proliferation, as revealed by Ki67 staining. *MYC* amplification is seen in ∼40% of HGSOC, and often co-occurs with *Brca1* alterations (Figure 1A). Retroviral over-expression of *Myc* in *Tp53*^*-/-*^*/Brca1*^*-/-*^ organoids (Figure 1B and C) resulted in additional increases in proliferation and organoid size, while impeding ciliary differentiation (Figure 1D and E). Orthotopic injection of *Tp53*^*-/-*^ or *Tp53*^*-/-*^*/Brca1*^*-/-*^ FTE cells (2 x 10^6^) failed to evoke tumors within a 6-month observation period. By contrast, mice injected with *Tp53*^*-/-*^;*Brca1*^*-/-*^;*Myc*^*OE*^ organoid cells developed ovarian tumors and omental metastases (Figure 1F), which resulted in the death of all mice within 4 months (Figure 1G). These tumors expressed markers associated with HGSOC, including PAX8, Cytokeratin 7 (CK7), P16, and Wilms Tumor 1 (WT1), and were strongly Ki67+ (Figure 1H). Hence, whereas compound BRCA1/TP53 deficiency is not sufficient to cause HGSOC, superimposing high MYC expression (or *Pten* and/or *Nf1* deletion; Table S1) results in a highly invasive, metastatic, lethal malignancy.

**Figure 1.**
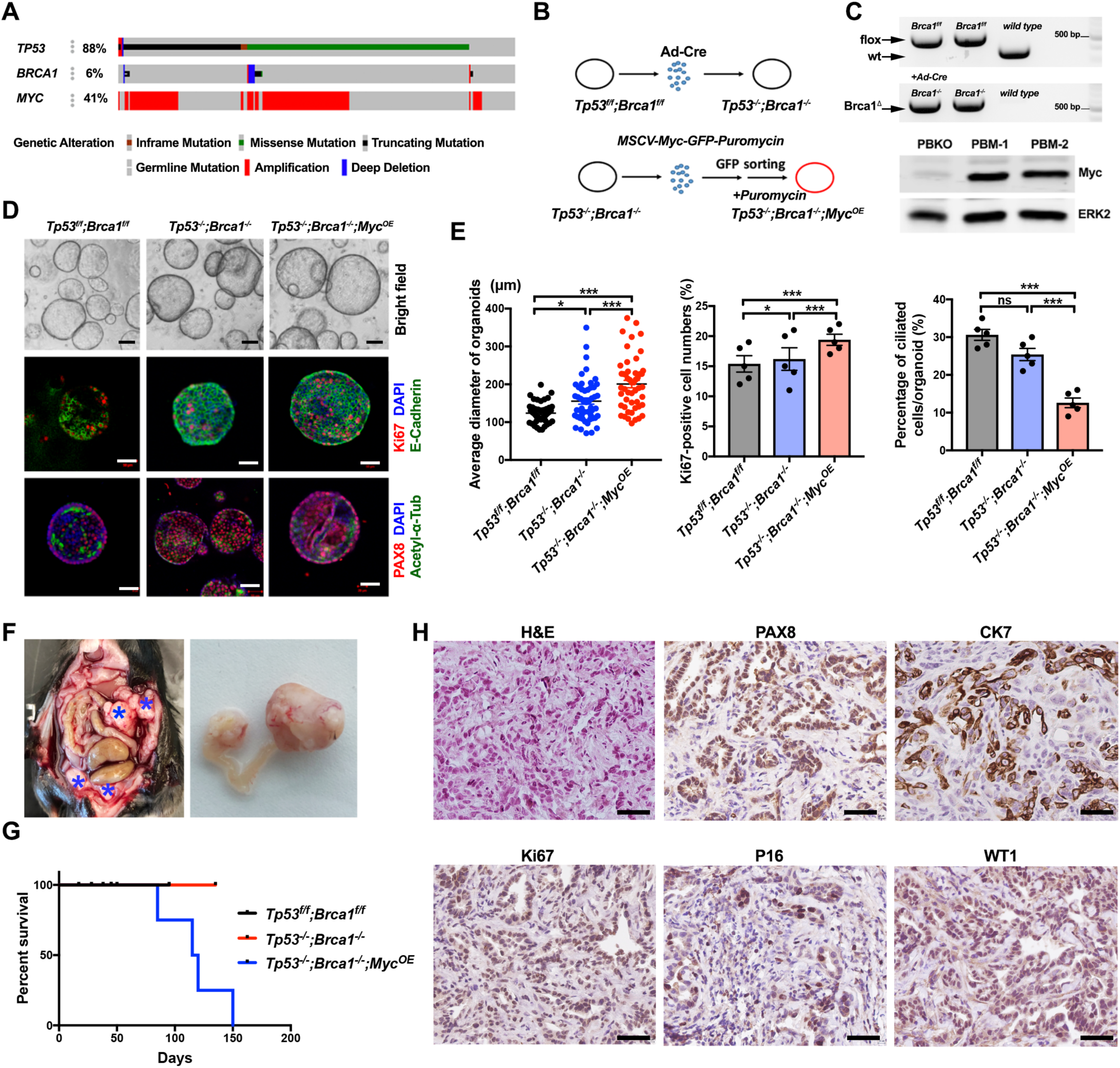
Generation of *Tp53*^*-/-*^;*Brca1*^*-/-*^;*Myc*^*OE*^ tumorigenic organoids. (A) OncoPrint showing genetic alterations of *TP53, BRCA1*, and *MYC* in human HGSOC (TCGA, Firehose Legacy). (B) Schematic showing approach used to generate *Tp53*^*-/-*^;*Brca1*^*-/-*^ (top panel) and *Tp53*^*-/-*^;*Brca1*^*-/-*^;*Myc*^*OE*^ (bottom panel) fallopian tube organoids from parental *Tp53*^*f/f*^;*Brca1*^*f/f*^ organoids. (C) Top panel: PCR of genomic DNA from the indicated organoids, Bottom panel: Representative MYC immunoblot showing two positive organoid clones. PBKO: *Tp53*^*-/-*^;*Brca1*^*-/-*^; PBM: *Tp53*^*-/-*^;*Brca1*^*-/-*^;*Myc*^*OE*^. (D) Representative bright field images of organoids and immunofluorescence staining with the indicated antibodies after 7 days in culture. Scale bars: 20 µm. (E) Average diameter of organoids (left panel), % Ki67-positive cells (middle panel) and % ciliated cells (right panel) from organoids of the indicated genotypes. Data represent mean ± SEM. ns, not significant, *P < 0.05, **P<0.01, ***P < 0.001, Tukey’s multiple comparison test. (F) Exposed abdomen of mouse 2 months after orthotopic injection of 2×10^6^ *Tp53*^*-/-*^;*Brca1*^*-/-*^;*Myc*^*OE*^ cells (left panel); asterisks indicate large metastatic deposits. Right panel: genital tract dissected from mouse at left showing large ovarian tumors (right panel). (G) K-M curves of mice following orthotopic injection of 2×10^6^ organoid cells of the indicated genotypes. n=6/group. (H) H&E-stained sections and immunohistochemical analysis of the indicated HGSOC markers in representative sections from *Tp53*^*-/-*^;*Brca1*^*-/-*^;*Myc*^*OE*^ ovarian tumors. Scale bars: 50 µm. See also Figure S1.

### *Tp53*^*-/-*^;*Pten*^*-/-*^;*Nf1*^*-/-*^ FTE organoids also cause HGSOC-like tumors

NF1 deficiency (due to inactivating mutation/deletion in *NF1*) is seen in ∼12% of HGSOC cases (Figure 2A) (Hollis and Gourley, 2016). *PTEN* loss also occurs fairly frequently (∼7%), and is associated with poor prognosis (Martins et al., 2014). We therefore assessed the effects of PTEN, NF1, or compound PTEN/NF1 deficiency on *Tp53*^*-/-*^ FTE (Figure 2B). We first used a lentiviral vector to introduce an sgRNA targeting *Pten* exon 2 into *Tp53*^*-/-*^ organoids. Three clones with bi-allelic deletion were identified and expanded, and PTEN deficiency was confirmed by immunoblotting (Figure 2C). An analogous strategy was used to target *Nf1* exon2 in *Tp53*^*-/-*^ or *Tp53*^*-/-*^;*Pten*^*-/-*^ organoids (Figure 2B and 2C). *Pten* deletion caused increased proliferation and organoid size, filling of the organoid lumen, and decreased ciliated cell differentiation. Although *Nf1* deletion (in *Tp53*^*-/-*^ FTE) decreased ciliary differentiation and altered organoid shape, proliferation and luminal integrity were unaltered. Triple-deleted (*Tp53*^*-/-*^; *Pten*^*-/-*^; *Nf1*^*-/-*^) and *Tp53*^*-/-*^;*Pten*^*-/-*^ organoids behaved similarly in these assays (Figure 2D and 2E).

**Figure 2.**
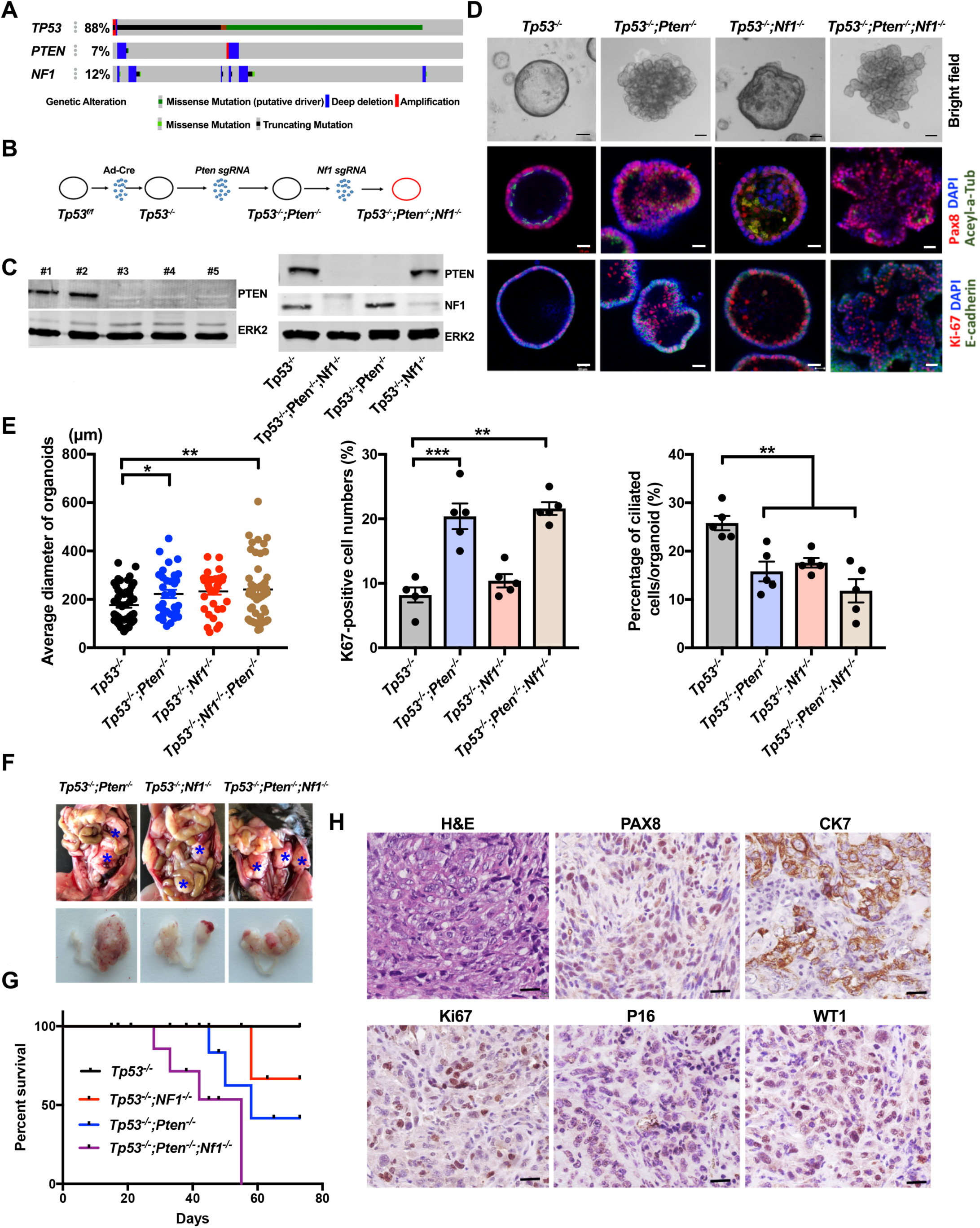
Generation of tumorigenic *Tp53*^*-/-*^;*Pten*^*-/-*^;*Nf1*^*-/-*^ organoids. (A) OncoPrint showing genetic alterations of *TP53, PTEN*, and *NF1* in human HGSOC (TCGA, Firehose Legacy). (B) Schematic showing approach used to generate *Tp53*^*-/-*^;*Pten*^*-/-*^ or *Tp53*^*-/-*^;*Pten*^*-/-*^;*Nf1*^*-/-*^ fallopian tube organoids from parental *Tp53*^*-/-*^ organoids. (C) Representative immunoblots for PTEN or NF1; ERK2 serves as a loading control. (D) Representative bright field organoid images and immunofluorescence staining for the indicated antibodies after 7 days in culture. Scale bars: 20 µm. (E) Average organoid diameter (left panel), % Ki67-positive cells (middle panel), and % ciliated cells (right panel) in organoids of the indicated genotypes. Data represent mean ± SEM. *P < 0.05, **P<0.01, ***P < 0.001, Tukey’s multiple comparison test. (F) Top panels: exposed abdominal cavities of mice bearing tumors of the indicated genotypes, 1 month after orthotopic injection of 2×10^6^ organoid cells of the indicated genotypes. Note the abdominal distention caused by massive ascites, large ovarian tumors, and peritoneal studding (asterisks). Bottom panel: genital tracts of mice in top panels. (G) K-M curves of mice following orthotopic injection with organoid cells (2×10^6^) of the indicated genotypes, n=6/group. (H) H&E-staining and IHC for the indicated HGSOC markers in representative sections from *Tp53*^*-/-*^;*Pten*^*-/-*^;*Nf1*^*-/-*^ tumors. Scale bars: 50 µm. Also see Figure S1.

We also tested the tumorigenicity of *Tp53*^*-/-*^;*Pten*^*-/-*^, *Tp53*^*-/-*^;*Nf1*^*-/-*.^, and *Tp53*^*-/-*^;*Pten*^*-/-*^;*Nf1*^*-/-*^ organoid cells (at least 2 clones each). Some double mutant-injected mice (8/20 *Tp53*^*-/-*^;*Pten*^*-/-*^, 8/24 *Tp53*^*-/-*^;*Nf1*^*-/-*^) developed tumors within 6 months of observation, but *Tp53*^*-/-*^;*Pten*^*-/-*^;*Nf1*^*-/-*^ cells showed more rapid and penetrant (28/30) tumorigenesis (Table S1). *Tp53*^*-/-*^;*Pten*^*-/-*^;*Nf1* cells also caused tumors more rapidly than did *Tp53*^*-/-*^;*Brca1*^*-/-*^;*Myc*^*OE*^ cells (compare Figure 1G and Figure 2G). *Tp53*^*-/-*^;*Pten*^*-/-*^;*Nf1*^*-/-*^ tumors metastasized to the omentum and evoked more massive ascites than did *Tp53*^*-/-*^;*Pten*^*-/-*^, *Tp53*^*-/-*^;*Nf1*^*-/-*^, *Tp53*^*-/-*^;*Brca1*^*-/-*^;*Myc*^*OE*^, or *Ccne1*^*OE*^ tumors (Figure 2F). Mice bearing *Tp53*^*-/-*^;*Pten*^*-/-*^;*Nf1*^*-/-*^ tumors also had shorter life spans, compared with each double knockout (Figure 2G). These tumors also expressed HGSOC-related markers (Figure 2H).

### *AKT2* and/or *KRAS* cooperate with *CCNE1* to cause HGSOC

Amplification or gain of *CCNE1*, which encodes the cell-cycle regulator CYCLIN E1, is the best characterized driver of HR-proficient HGSOC, accounting for ∼20% of HGSOC overall (Konstantinopoulos et al., 2015; Nakayama et al., 2010). *AKT2* and *KRAS* amplification occur in 8% and 16%, respectively, of HGSOC cases, and co-occur with *CCNE1* amplification (Figure 3A). To model *CCNE1* amplification alone or in concert with *KRAS* and/or *AKT2, Ccne1, Akt2*, and/or *Kras* were over-expressed sequentially in *Tp53*^*-/-*^ FTE by transduction with lentiviruses harboring different selection markers (neomycin, blasticidin, mCherry). Over-expression of each protein was confirmed by immunoblotting (Figure 3B and Figure S2B). Organoid diameter and gross morphology were not affected significantly by CCNE1, AKT2, or KRAS over-expression alone (compared with parental *Tp53*^*-/-*^ organoids). Overexpression of CCNE1, but not AKT2 or KRAS, significantly increased proliferation (Figure 3B). Most likely, this increase was offset by a comparable increase in cell death, accounting for the lack of alteration of organoid size; *CCND1* over-expression has analogous effects on MCF10A mammary organoids (Debnath et al., 2002). Superimposing *Akt2* or *Kras* overexpression on *Tp53*^*-/-*^;*Ccne1*^*OE*^ organoids resulted in further enhanced proliferation, increased organoid diameter, luminal filling, and organoid disorganization, which was even more pronounced in quadruple mutants. *Ccne1* over-expression alone did not alter ciliated cell differentiation, but ciliated cells were virtually undetectable in triple and quadruple mutant organoids (Figures 3C and 3D). *Tp53*^*-/-*^;*Ccne1*^*OE*^ FTE cells did not give rise to tumors, even at 6 months after orthotopic injection. By contrast, *Tp53*^*-/-*^;*Ccne1*^*OE*^;*Akt2*^*OE*^, *Tp53*^*-*^ */-;Ccne1*^*OE*^;*Kras*^*OE*^ and *Tp53*^*-/-*^;*Ccne1*^*OE*^;*Akt2*^*OE*^;*Kras*^*OE*^ formed large, palpable ovarian tumor masses, accompanied by massive omental metastasis and death within 2 months following injection (Figure 3E). There was no apparent difference in tumor formation between mice bearing each triple mutant, but quadruple mutants showed accelerated tumorigenesis (Figure 3F). *Tp53*^*-*^ */-;Ccne1*^*OE*^;*Akt2*^*OE*^;*Kras*^*OE*^ tumors displayed features of high grade, poorly differentiated, invasive carcinoma expressing HGSOC markers (Figure 3G).

**Figure 3.**
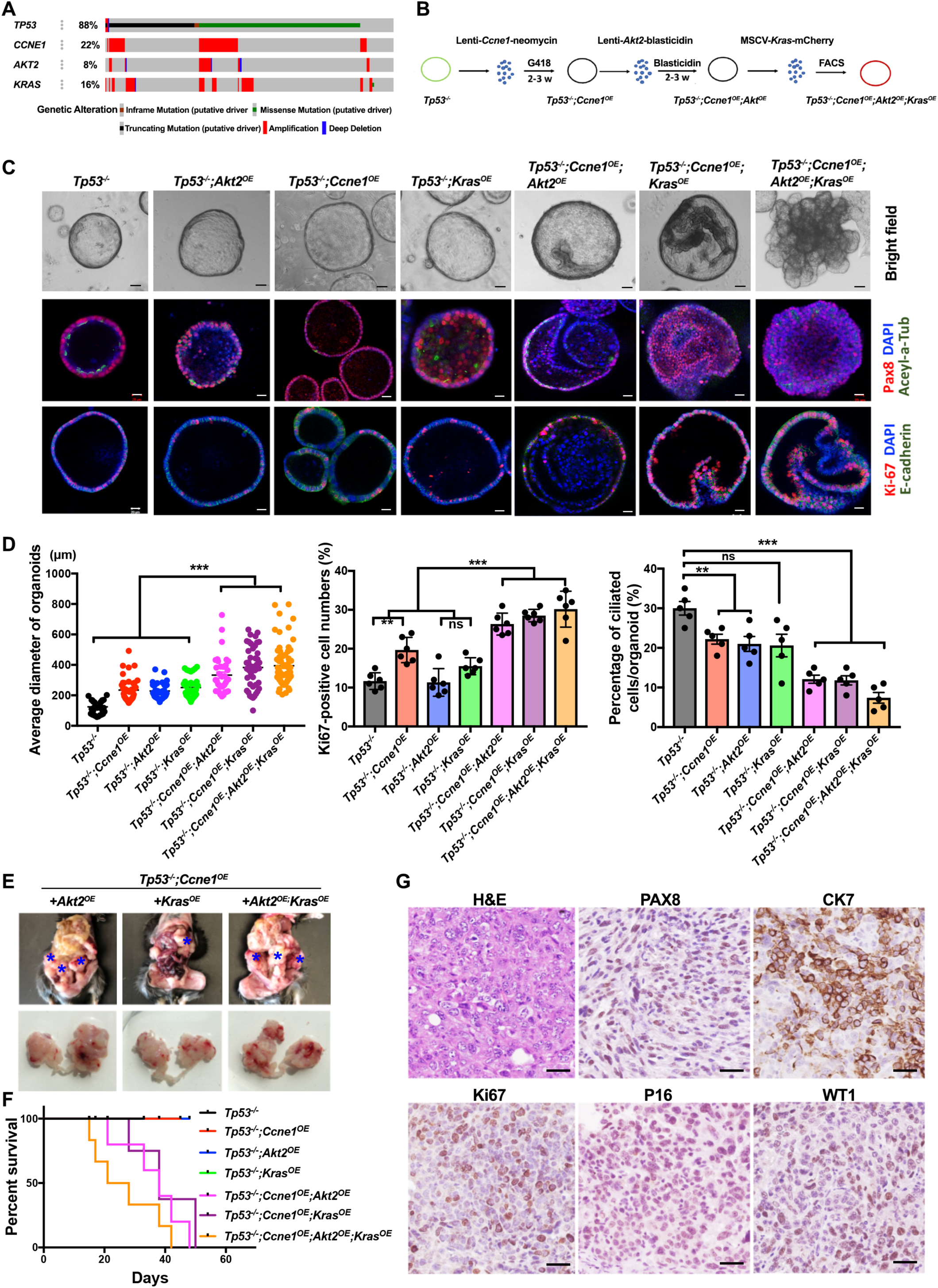
*Akt2 and/*or *Kras* over-expression in *Tp53*^*-/-*^;*Ccne1*^*OE*^ organoids causes HGSOC-like tumors. (A) OncoPrint showing genetic alterations of *Tp53, CCNE1, AKT2* and *KRAS* in human HGSOC (TCGA, Firehose Legacy). (B) Schematic showing generation of the indicated organoids, starting from *Tp53*^*-/-*^ fallopian tube cells. (C) Representative bright field and immunofluorescence micrographs of organoids with the indicated genotypes. Scale bars: 20 µm. (D) Average organoid diameter (left panel), % Ki67-positive cells (middle panel), and % ciliated cells (right panel) in organoids with the indicated genotypes. Data represent mean ± SEM. *P < 0.05, **P<0.01, ***P < 0.001, Tukey’s multiple comparison test. (E) Exposed abdomens of mice 1 month after orthotopic injection of 2×10^6^ organoid cells of the indicated genotypes. Asterisks show representative tumor deposits. Bottom panel: genital tracts of mice in top panels. (F) K-M curves of mice injected with 2×10^6^ organoid cells from the indicated genotypes. n=6/group. (G) H&E-stained sections and IHC for the indicated HGSOC markers in tumor from representative *Tp53*^*-/-*^;*Ccne1*^*OE*^;*Akt2*^*OE*^;*Kras*^*OE*^ mouse. Scale bars: 50 µm. See also Figure S1 and Figure S2

Taken together, the above data establish that several different combinations of genetic abnormalities seen in human HGSOC give rise to lethal ovarian cancers in immune-competent mice, and begin to assign complementation groups for various *in vitro* properties (proliferation, differentiation, organoid morphology) and tumorigenic capacity. Other combinations of genetic abnormalities reported in TCGA also give rise to HGSOC in mice (Table S1).

### Ovarian tumor genotype dictates tumor microenvironment

We suspected that different HGSOC genotypes might elicit distinct TMEs. To test this hypothesis, we assayed *Tp53*^*-/-*^;*Brca1*^*-/-*^;*Myc*^*OE*^, *Tp53*^*-/-*^;*Pten*^*-/-*^;*Nf1*^*-/-*^, and *Tp53*^*-/-*^;*Ccne1*^*OE*^;*Akt2*^*OE*^;*Kras*^*OE*^ tumors by flow cytometry using lymphoid and myeloid marker panels (see Figure S3A and S3B for markers and gating). The percentage of CD45^+^ immune cells (compared with CD45^-^ tumor/stromal cells) was significantly higher in *Tp53*^*-/-*^;*Ccne1*^*OE*^;*Akt2*^*OE*^;*Kras*^*OE*^ and *Tp53*^*-/-*^;*Brca1*^*-/-*^;*Myc*^*OE*^ tumors, compared with *Tp53*^*-/-*^;*Pten*^*-/-*^;*Nf1*^*-/-*^ tumors (Figure S3C). None of the tumors had many B (CD19^+^), NK (NK1.1^+^), or NKT (NK1.1^+^CD3^+^) cells (Figure 4A and Figure S3C). Nevertheless, the cellular composition of the CD45^+^ population in tumors of different genotype differed substantially (Figure 4A and 4B).

**Figure 4.**
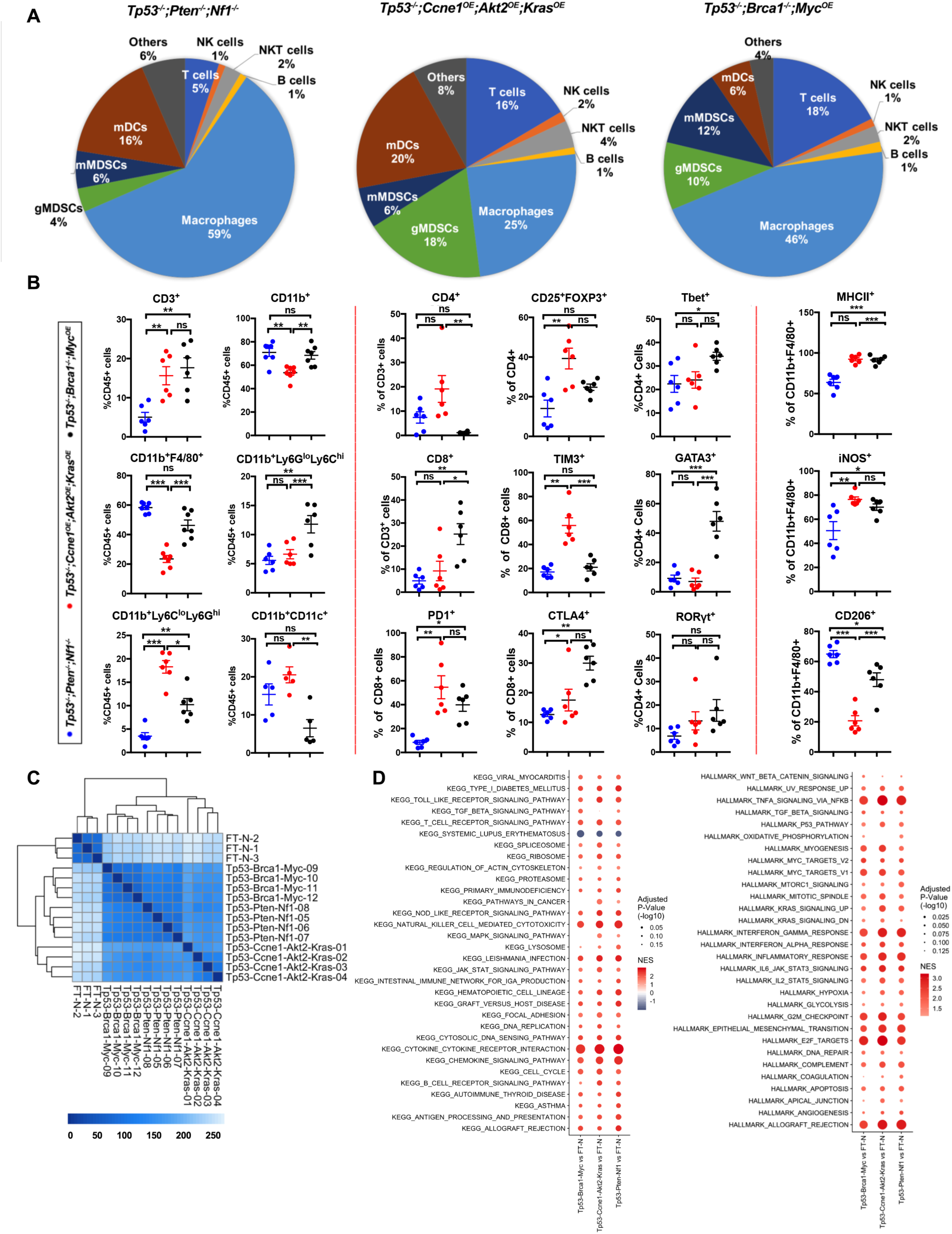
Tumor genotype determines immune landscape and transcriptome. (A) Pie charts summarizing composition of immune cells (CD45+) in tumors with the indicated genotypes. Note that CD45+ cells (as a % of total tumor cells) were significantly less in *Tp53*^*-/-*^;*Pten*^*-/-*^;*Nf1*^*-/-*^ tumors, but similar in the other two genotypes (see Supplementary Figure 3). (B) Immune cell subtyping by flow cytometric analysis of representative tumors of the indicated genotypes. Each point represents a tumor from a different mouse. Data are presented as mean ± SEM, **P<0.01, ***P<0.001, 2-way ANOVA. (C) Heat map showing sample distances by hierarchical clustering, based on variance stabilized expression levels of all genes in normal FT, *Tp53*^*-/-*^;*Ccne1*^*OE*^;*Akt2*^*OE*^;*Kras*^*OE*^, *Tp53*^*-/-*^;*Pten*^*-/-*^;*Nf1*^*-/-*^ and *Tp53*^*-/-*^;*Brca1*^*-/-*^;*Myc*^*OE*^ tumors, respectively. Shading represents Euclidian distance for each sample pair. (D) Pathway analysis comparing normal FT and cognate tumors of the indicated genotypes. Significantly enriched KEGG (left) and MSigDB Hallmark genes (right) of the genes ranked by fold change between the indicated groups are shown. Size of each circle represents the log-transformed FDR-adjusted p-value within each category; color indicates normalized enrichment score. Also see Figure S4A.

*Tp53*^*-/-*^;*Pten*^*-/-*^;*Nf1*^*-/-*^ tumors had a predominant macrophage population (CD11b^+^F4/80^+^), smaller populations of myeloid dendritic cells (mDC, CD11b^+^CD11C^+^), granulocytic myeloid-derived suppressor cells (g-MDSC, CD11b^+^Ly6C^lo^Ly6G^hi^), and monocytic myeloid-derived suppressor cells (m-MDSC, CD11b^+^Ly6G^lo^Ly6C^hi^), and sparse T lymphocytes (CD3^+^). As these tumors had fewer CD45+ cells overall (Figure S3C), the absolute number of T cells in *Tp53*^*-/-*^;*Pten*^*-/-*^;*Nf1*^*-/-*^ tumors is even lower, compared with the other models. The macrophages in *Tp53*^*-/-*^;*Pten*^*-/-*^;*Nf1*^*-/-*^ tumors had greater “M2-like” character, with high percentages of CD11b^+^ F4/80^+^ cells expressing CD206 and a lower percentage of iNOS^+^ cells. Most macrophages in these tumors co-expressed M1 and M2 markers, though, more consistent with an “M0-like” state (Murray et al., 2014; Reinartz et al., 2014).

*Tp53*^*-/-*^;*Ccne1*^*OE*^;*Akt2*^*OE*^;*Kras*^*OE*^ tumors were more inflamed, exhibiting infiltration with macrophages, mDCs, g-MDSCs, and T lymphocytes (Figure 4A and 4B). However, nearly half of the CD4^+^ T cells in these tumors were T regulatory cells (Tregs, CD25^+^Foxp3^+^), while most of the CD8^+^ cells expressed markers of “exhaustion” (TIM3+, PD1+). The macrophage population (F4/80^+^) in these mice expressed “M1-like” (MHCII^+^, iNOS^+^) and “M2-like” (CD206^+^) cells, although the former predominated.

Finally, *Tp53*^*-/-*^;*Brca1*^*-/-*^;*Myc*^*OE*^ tumors had large percentages of macrophages and lower fractions of g-MDSCs, m-MDSCs, and mDCs (Figure 4A and 4B). In contrast to the other models, the CD4^+^ and CD8^+^ T cells in *Tp53*^*-/-*^;*Brca1*^*-/-*^;*Myc*^*OE*^ tumors were predominantly (>60%) CD44^+^ and strongly CTLA4^+^ and PD1^+^ (Figure 4B and Figure S3C), consistent with prior activation. Moreover, compared with the cognate cells in *Tp53*^*-/-*^;*Ccne1*^*OE*^;*Akt2*^*OE*^;*Kras*^*OE*^ tumors, CD8^+^ cells in *Tp53*^*-/-*^;*Brca1*^*-/-*^;*Myc*^*OE*^ tumors had lower levels of TIM3-positivity, comporting with less exhaustion, and there were fewer Treg cells. *Tp53*^*-/-*^;*Brca1*^*-/-*^;*Myc*^*OE*^ tumors had more balanced populations of Th1 and Th2 cells (Th1/Th2: 0.7), whereas the other models mostly had Th1 cells (Th1/Th2: 2.4 in *Tp53*^*-/-*^;*Pten*^*-/-*^;*Nf1*^*-/-*^; Th1/Th2: 3.4 in *Tp53*^*-/-*^ in *Ccne1*^*OE*^;*Akt2*^*OE*^;*Kras*^*OE*^), and *Tp53*^*-/-*^;*Brca1*^*-/-*^;*Myc*^*OE*^ macrophages showed more M1-like character (%CD206/%iNos:1.3) than did the other models (%CD206/%iNos: 0.2 in *Ccne1*^*OE*^;*Akt2*^*OE*^;*Kras*^*OE*^; %CD206/%iNos: 0.6 in *Tp53*^*-/-*^;*Brca1*^*-/-*^;*Myc*^*OE*^).

PD-L1 was also expressed variably on tumor cells and on different cell types in the TME in a genotype-dependent manner. In all models, approximately 40-45% of m-MDSCs were PD-L1+. In *Ccne1*^*OE*^;*Akt2*^*OE*^;*Kras*^*OE*^ tumors, 45% of g-MDSCs also expressed PD-L1, whereas expression on gMDSCs was lower in the *Tp53*^*-/-*^;*Brca1*^*-/-*^;*Myc*^*OE*^ (25%) and *Tp53*^*-/-*^;*Pten*^*-/-*^;*Nf1*^*-/-*^ (14%) models. By contrast, 58% of the macrophages in *Tp53*^*-/-*^;*Pten*^*-/-*^;*Nf1*^*-/-*^ tumors were PD-L1+ *Tp53*^*-/-*^;*Pten*^*-/-*^;*Nf1*^*-/-*^, *Ccne1*^*OE*^;*Akt2*^*OE*^;*Kras*^*OE*^, and *Tp53*^*-/-*^;*Brca1*^*-/-*^;*Myc*^*OE*^ tumors showed PD-L1 expression on 2%, 8%, and 5% expression on CD45-cells (malignant cells, respectively (data not shown). The distinct TMEs of evoked by tumors of different genotypes argue against the utility of using a single “HGSOC” model, such as ID8 cells (much less one not bearing the canonical mutations associated with the disease), for testing immune therapies for this disease.

### Ovarian tumors with different genotypes have distinct transcriptomes

We used RNA sequencing to analyze the transcriptomes of tumors (4 each) of each genotype and normal FT. Unsupervised hierarchical clustering and principle components analysis (PCA) revealed clear separation between tumor and normal samples (Figure 4C). Each tumor model also had a distinct transcriptome by unsupervised clustering, with the *Tp53*^*-/-*^;*Ccne1*^*OE*^;*Akt2*^*OE*^;*Kras*^*OE*^ model showing the greatest difference from the other two. Pathway analysis revealed that, compared with normal FT, tumor transcriptomes were enriched primarily for KEGG gene sets associated with the immune response (e.g., cytokine/cytokine receptor interaction, chemokine signaling pathway, antigen processing and presentation, Leishmania infection, Toll like receptor signaling pathway, etc.), and, to a lesser extent, for processes related to proliferation (e.g., DNA replication, cell cycle, ribosome, etc.). Similar enrichment was seen for Hallmark Gene sets associated with inflammatory/immune (allograft rejection, TNFα signaling, nterferon gamma response, interferon alpha response, complement signaling, etc.) and proliferative (G2/M checkpoint, MYC targets, KRAS signaling, mTORC signaling, etc.) processes. Tumor samples also were enriched for multiple Oncogenic Gene Sets (Figure 4D and Figure S4A).

Pairwise comparisons also revealed significant differences, several that comported with their distinct genotypes. For example, compared with *Tp53*^*-/-*^;*Ccne1*^*OE*^;*Akt2*^*OE*^;*Kras*^*OE*^ tumors, the *Tp53*^*-/-*^;*Pten*^*-/-*^;*Nf1*^*-/-*^ model showed lower expression of “Pten-down” genes and of “MEK-up,” “KRAS-up” and “EGFR-up” genes; these findings likely reflect stronger activation of the RAS/ERK pathway in KRAS over-expressing, compared with NF1-deficient, tumor cells. By contrast, “KRAS-up” gene sets were enriched in *Tp53*^*-/-*^;*Pten*^*-/-*^;*Nf1*^*-/-*^ tumors compared with their *Tp53*^*-/-*^;*Brca1*^*-/-*^;*Myc*^*OE*^ counterparts, and also showed higher levels of “AKT-up” gene sets (Figure S4A).

To begin to understand the molecular basis for their distinct TMEs, we compared chemokine, cytokine, and hematopoietic growth factor gene expression in each type of tumor (Figure 5A). Most interleukins were expressed at low/undetectable levels in all models, as were many chemokines. IL15, IL16, IL18, IL33, and IL34 were expressed significantly in all tumors. *Lif, IL1b, Csf1* (MCSF) and to a lesser extent, *Tnf*α, were expressed at higher levels in *Tp53*^*-/-*^;*Pten*^*-/-*^;*Nf1*^*-/-*^ and *Tp53*^*-/-*^;*Ccne1*^*OE*^; *Akt2*^*OE*^, compared with *Tp53*^*-/-*^;*Brca1*^*-/-*^;*Myc*^*OE*^, tumors. Some chemokines (e.g., *Cxcl12, Cxcl16*) were expressed at similar, high levels in all models (Figure 5A). Others were expressed differentially depending on tumor genotype: for example, *Ccl2* and *Ccl5* were expressed most highly in *Tp53*^*-/-*^;*Ccne1*^*OE*^;*Akt2*^*OE*^;*Kras*^*OE*^ tumors, at intermediate levels in *Tp53*^*-/-*^;*Pten*^*-/-*^;*Nf1*^*-/-*^ tumors, and at lower levels in *Tp53*^*-/-*^;*Brca1*^*-/-*^;*Myc*^*OE*^ tumors. *Cxcl1* levels were higher in *Tp53*^*-/-*^;*Brca1*^*-/-*^;*Myc*^*OE*^ tumors and *Tp53*^*-/-*^;*Pten*^*-/-*^;*Nf1*^*-/-*^ tumors. *Ccl6-9* were expressed in most tumors, although at generally lower levels in *Tp53*^*-/-*^;*Brca1*^*-/-*^;*Myc*^*OE*^ tumors. By contrast, *Cxcl9* was expressed at highest levels in *Tp53*^*-/-*^;*Brca1*^*-/-*^;*Myc*^*OE*^ tumors and at lowest levels in *Tp53*^*-/-*^;*Ccne1*^*OE*^; *Akt2*^*OE*^, *Kras*^*OE*^ tumors, whereas *Cxcl10* was expressed at higher levels in the latter. Notably, *Vegfa* and *Tgfb1* were expressed at high levels in all tumors (Figure 5A).

**Figure 5.**
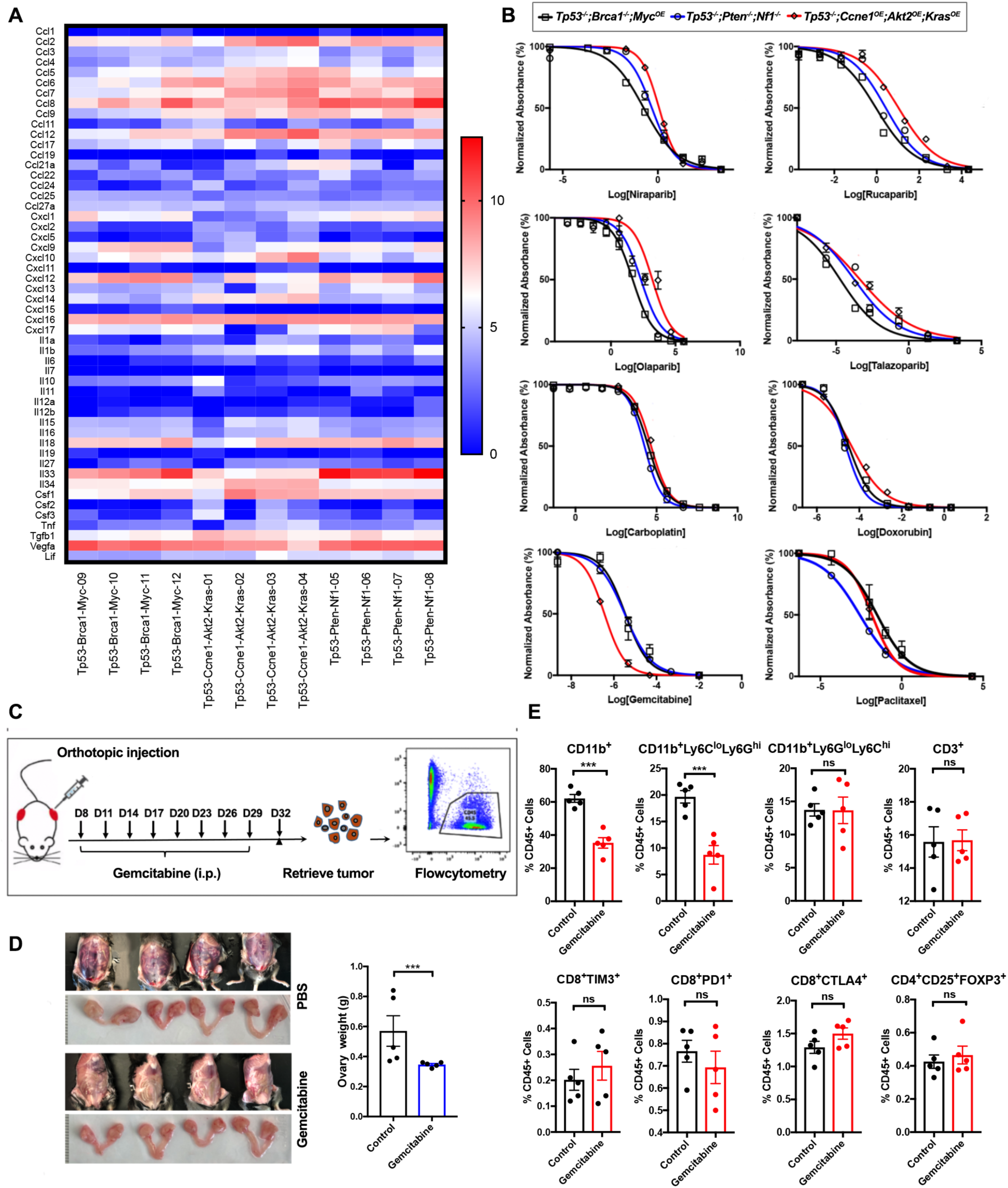
Tumor genotype causes distinct patterns of cytokine gene expression and drug sensitivity. (A) Cytokine/chemokine transcript levels (Log-transformed TPMs) of the indicated chemokines/cytokines in representative tumors from each genotype. (B) Dose-response curves for the indicated drugs in tumorigenic organoid lines with different genotypes. Cell viability was calculated relative to 0.01% DMSO-treated control cells, measured after 5 days of treatment. (C) Schematic depicting gemcitabine treatment of *Tp53*^*-/-*^;*Ccne1*^*OE*^;*Akt2*^*OE*^;*Kras*^*OE*^ tumor-bearing mice. (D) Exposed abdominal cavities (Top and third panels) and genital tracts (second and fourth panels) of mice treated with PBS (upper two panels) or with Gemcitabine (lower two panels), showing abdominal distention due to ascites and larger ovarian tumors in the PBS group. Right panel: Ovary weight in PBS- and gemcitabine-treatment groups at the end of treatment. (E) Flow cytometric analysis for the indicated immune subsets in *Tp53*^*-/-*^;*Ccne1*^*OE*^;*Akt2*^*OE*^;*Kras*^*OE*^ tumors with or without Gemcitabine treatment. Data indicate means ± SEM, ns, not significant, ***p < 0.001, unpaired t test. See also Figure S4.

These complex, yet distinct, mixes of immune modulatory factors, acting in concert, presumably sculpt the microenvironments of these tumors. While it is difficult to attribute with certainty differences in specific TME components to particular factors, several specific, testable hypotheses emerge. For example, the T cell infiltrate in *Tp53*^*-/-*^;*Ccne1*^*OE*^;*Akt2*^*OE*^;*Kras*^*OE*^ tumors is likely to be directed by CXCL10 with a lesser contribution of CXCL9, whereas the latter is probably most responsible for T cell infiltration in *Tp53*^*-/-*^;*Brca1*^*-/-*^;*Myc*^*OE*^ tumors. Similarly, whereas CCL2 and MCSF probably contribute to the macrophage infiltration in *Tp53*^*-/-*^;*Pten*^*-/-*^;*Nf1*^*-/-*^ and *Tp53*^*-/-*^;*Ccne1*^*OE*^;*Akt2*^*OE*^;*Kras*^*OE*^ tumors (with potential contributions from CCL6-9), *Tp53*^*-/-*^;*Brca1*^*-/-*^;*Myc*^*OE*^ -associated macrophages might be driven primarily by CXCL1 and CCL6-9.

Ultimately, the TME results from the combined effects of tumor-derived factors, which drive initial immune cell influx, interacting with factors produced by the infiltrating immune cells. To begin to deconvolve this complexity, we assessed the “secretome” of each type of tumorigenic organoid (using Luminex® assays). Like their cognate tumors, organoids secreted a complex, genotype-specific mix of factors (Figure S4B). Comparing these factors to the tumor transcriptome suggests that many tumor-derived cytokines/chemokines initiate and help to maintain the TME (e.g., CCL2, CCL5, CXCL10 for *Tp53*^*-/-*^;*Ccne1*^*OE*^;*Akt2*^*OE*^;*Kras*^*OE*^ tumors; MCSF, CXCL1, and CXCL9 for *Tp53*^*-/-*^;*Brca1*^*-/-*^;*Myc*^*OE*^ tumors; MCSF, CXCL1, CCL2, and VEGF-A for *Tp53*^*-/-*^;*Pten*^*-*^ */-;Nf1*^*-/-*^ tumors). CCL2, CCL5 and CXCL10 were also expressed at high levels in the serum of tumor-bearing *Tp53*^*-/-*^;*Ccne1*^*OE*^;*Akt2*^*OE*^;*Kras*^*OE*^ mice (data not shown). Other factors might contribute to TME initiation but are no longer expressed at high levels in tumors themselves (e.g., G-CSF/*Csf2* in *Tp53*^*-/-*^;*Brca1*^*-/-*^;*Myc*^*OE*^ and *Tp53*^*-/-*^;*Pten*^*-/-*^;*Nf1*^*-/-*^ tumors). Some presumably emanate from primarily from tumor-infiltrating immune cells, rather than malignant epithelial cells themselves (e.g., GMSCF/*Csf3*, CXCL5 in *Tp53*^*-/-*^;*Brca1*^*-/-*^;*Myc*^*OE*^, among others).

### Organoid genotype alters drug sensitivity

We tested these models for sensitivity to several FDA-approved drugs for HGSOC (Figure 5B) and investigational/experimental agents (Figure S4C). Organoids were titurated into small clumps, dissociated into single cells, and dispensed into 96-well Matrigel pre-coated plates (see Experimental Procedures). Each agent was added at various doses, and cell viability was assessed 5 days later. As expected, *Tp53*^*-/-*^;*Brca1*^*-/-*^;*Myc*^*OE*^ cells showed increased sensitivity to PARP-Is (Figure 5B), although differential sensitivity varied for individual PARP-Is and was less than with conventional ovarian cancer cell lines (Baloch et al., 2019). *Brca1*-deleted cells showed slightly increased sensitivity to Carboplatin, although there was substantial overlap with the other mutants. Conceivably, MYC over-expression reduces PARP-I and/or platinum sensitivity in *Brca1*^*-/-*^ cells (Carey et al., 2018; Reyes-Gonzalez et al., 2015). *Tp53*^*-/-*^;*Ccne1*^*OE*^;*Akt2*^*OE*^;*Kras*^*OE*^ organoids were more sensitive to gemcitabine than the other models, consistent with the increased replication stress caused by *CCNE1* amplification (Hill et al., 2018). By contrast, and unexpectedly, *Tp53*^*-/-*^;*Pten*^*-/-*^;*Nf1*^*-/-*^ cells showed enhanced susceptibility to paclitaxel (Figure 5B). *Tp53*^*-/-*^;*Brca1*^*-/-*^;*Myc*^*OE*^ and *Tp53*^*-/-*^;*Pten*^*-/-*^;*Nf1*^*-/-*^ organoids had increased sensitivity to the ATR inhibitor BAY1895344, whereas chloroquine, which inhibits endosomal acidification and is often used as an autophagy inhibitor, was differentially toxic for all genotypes (*Tp53*^*-/-*^;*Pten*^*-/-*^;*Nf1*^*-/-*^>*Tp53*^*-/-*^;*Ccne1*^*OE*^;*Akt2*^*OE*^;*Kras>Tp53*^*-/-*^;*Brca1*^*-/-*^;*Myc*^*OE*^). All genotypes showed comparable sensitivity to the CDK7 inhibitor YKI-5-1241, the CDK7/9 inhibitor PHA767491, and CDK2/7/9 inhibitor Seliciclib (Figure S4C).

### Rationally derived combination therapy yields durable complete responses in *Tp53*^*-/-*^;*Ccne1*^*OE*^;*Akt*^*OE*^;*Kras*^*OE*^ HGSOC

We next sought to assess the utility of our tumorigenic organoid platform for developing therapies for HGSOC. To enable rapid clinical translation, we focused on approved drugs and on CCNE1-overexpressing tumors, given the particularly poor prognosis of this subgroup and their limited response to current therapies. Consistent with our *in vitro* findings, gemcitabine administration to mice with *Tp53*^*-/-*^;*Ccne1*^*OE*^;*Akt2*^*OE*^;*Kras*^*OE*^ tumors reduced disease burden, but there were no complete responses (CRs) (Figures 5C and 5D). As in other tumor models (Bezu et al., 2015; Deshmukh et al., 2018; Suzuki et al., 2005), gemcitabine treatment also reduced g-MDSCs (CD11b^+^Ly6C^lo^LY6G^hi^) in the TME, but other cell populations, most notably Tregs (CD24^+^CD25^+^FoxP3^+^) and T cells expressing exhaustion markers (TIM3/LAG3/PD1), remained unchanged (Figure 5E and data not shown).

Based on these data, we designed a regimen designed to attack tumor cells while normalizing the TME (Figure 6A): gemcitabine to cytoreduce tumor cells and g-MDSCs, anti-CTLA4 antibodies to target Tregs (Walker, 2013), and anti-PD-L1 antibodies to reactivate exhausted CD8 cells (Huang et al., 2017; Wei et al., 2017). This combination produced CRs in 10/10 treated mice, pooled from two experiments (Figure 6B and 6C, Figures S5A and S5B). Following treatment cessation (at day 35; Figure 6A), tumors failed to recur over a 60-day observation period (Figure 6D and Figure S5C). Gemcitabine plus anti-PD-L1 (but not anti-CTLA4) evoked a greater decrease in tumor burden and ascites than gemcitabine alone, but no CRs. Gemcitabine/anti-CTLA4 reduced ascites but did not measurably diminish tumor burden (Figure 6C, Figures S5A and S5B). Tumors recurred in all mice in the 2-drug combination groups after therapy was stopped, leading rapidly to death (Figure 6D). Histological analysis of mice sacrificed after 8 cycles of therapy revealed normal fat tissue abutting minimal amounts of residual tumor in the injected bursae of gemcitabine/anti-PD-L1/anti-CTLA4-treated animals; by contrast, considerable tumor remained in mice treated with gemcitabine/anti-PDL1 or gemcitabine/anti-CTLA4 (Figure 6E). Multi-color immunofluorescence (IF) confirmed that Ly6G^+^ cells were decreased in mice treated with gemcitabine, alone or in combination with anti-CTLA4 and/or anti-PDL1. Only tumors from mice treated with the triple combination showed significant increased T cell (CD3^+^) infiltration, which included CD4^+^ and CD8^+^ T cells (Figure 6F). Combination-treated mice also showed increases in granzyme B+ (cytolytic) cells, and decreased numbers of macrophages and Tregs (Figure S6).

**Figure 6.**
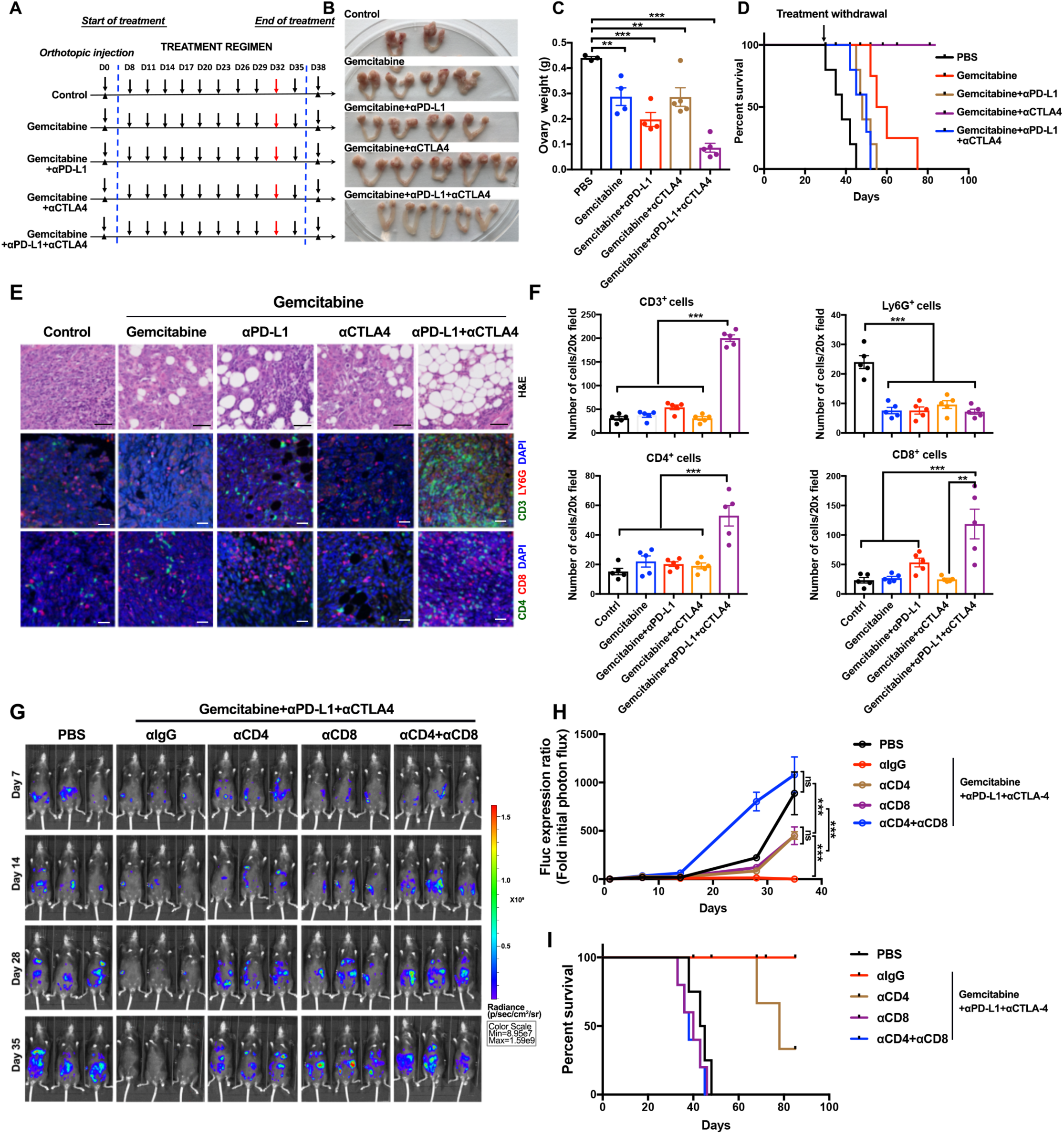
Rationally derived combination regimen results in complete responses in *Tp53*^*-/-*^;*Ccne1*^*OE*^;*Akt2*^*OE*^;*Kras*^*OE*^ tumors. (A) Schematic depicting treatment regimens (n=10 mice/group, 2 batches). For each aet of experiments, five mice were sacrificed at Day 32 for histological analysis; the other 5 were continued on treatment until Day 35, then treatment was withdrawn and mice were followed thereafter for survival. (B) Representative genital tracts from *Tp53*^*-/-*^;*Ccne1*^*OE*^;*Akt2*^*OE*^;*Kras*^*OE*^ tumor-bearing mice treated as indicated; mice were sacrificed at Day 32 of the scheme in (**A**). (C) Ovary weights from mice in the indicated treatment groups. Each point represents one mouse. (D) K-M curves of *Tp53*^*-/-*^;*Ccne1*^*OE*^;*Akt2*^*OE*^;*Kras*^*OE*^ tumor-bearing mice, treated as indicated in **A** until Day35, and then monitored for recurrence, n=5 mice/group. (E) H&E and IF staining for the indicated immune markers and DAPI (nuclei) in ovarian sections from the indicated groups. Note that the ovarian fat pad has almost no tumor after Gemcitabine+αPD-L1+αCTLA4 treatment. See also Supplementary Figure 6. Black scale bars: 50 µm, while scale bars: 20 µm. (F) Quantification of the indicated immune cells from the sections in (**E**). Each point represents average cell number per 20X field from 5 independent sections of each mouse. Error bars indicate SEM; **P<0.01, ***P<0.001, 2-way ANOVA. (G) Representative bioluminescence images of mice bearing orthotopic tumor allografts (expressing luciferase), treated as indicated, and measured at Days 7, 14, 28 and 35, respectively. (H) Relative photon flux, quantified by the intensity of bioluminescence in the regions of interest (ROIs), determined at the indicated times in mice from each treatment group, n=5 mice/group. Error bars indicate SEM; ns, not significant, ***P<0.001, 2-way ANOVA. (I) K-M curves for *Tp53*^*-/-*^;*Ccne1*^*OE*^;*Akt2*^*OE*^;*Kras*^*OE*^ tumor-bearing mice, treated as indicated. See also Figure S5 and Figure S6.

### Durable responses are T cell-dependent

The durability of the responses seen in gemcitabine/anti-CTLA4/anti-PD-L1-treated mice, and the attendant T cell influx, prompted us to ask if these responses were T cell-dependent. To this end, we depleted CD4+ and/or CD8+ T cells and re-assessed treatment efficacy. To enhance our ability to monitor tumors, we transduced *Tp53*^*-/-*^;*Ccne1*^*OE*^;*Akt2*^*OE*^;*Kras*^*OE*^ organoids with a luciferase-expressing lentivirus prior to implantation; preliminary experiments showed that luciferase-expressing tumors behaved similarly to the parental controls (Figure S5C and data not shown). Depletion of the expected T cell population was confirmed by flow cytometry of peripheral blood (Figure S5D and S5E). CD4 or CD8 cell depletion impaired the anti-tumor response of the combination regimen, whereas tumors from mice lacking CD4 and CD8 T cells actually grew faster in the presence of the combination therapy than did tumors in PBS-treated mice with intact immune systems (Figure 6G and H). Survival of combination-treated, CD8- or CD4+CD8-depleted tumor bearing mice was the same as PBS-treated tumor-bearing mice with intact immune systems, whereas CD4-depletion also impaired survival in the combination-treated group, but to a lesser extent.

### Therapeutic efficacy is tumor genotype-specific

To test whether the efficacy of gemcitabine/anti-CTLA4/anti-PD-L1 was specific for *Tp53*^*-/-*^;*Ccne1*^*OE*^;*Akt2*^*OE*^;*Kras*^*OE*^ tumor-bearing mice, we tested this regimen in mice with *Tp53*^*-*^ */-;Pten*^*-/-*^;*Nf1*^*-/-*^ tumors. Remarkably, the latter mice were completely refractory to this three-drug combination, as measured by tumor burden, percentage of mice with ascites, and survival (Figures 7A and 7C). Finally, we tested the effects of single agent paclitaxel on these models.

**Figure 7.**
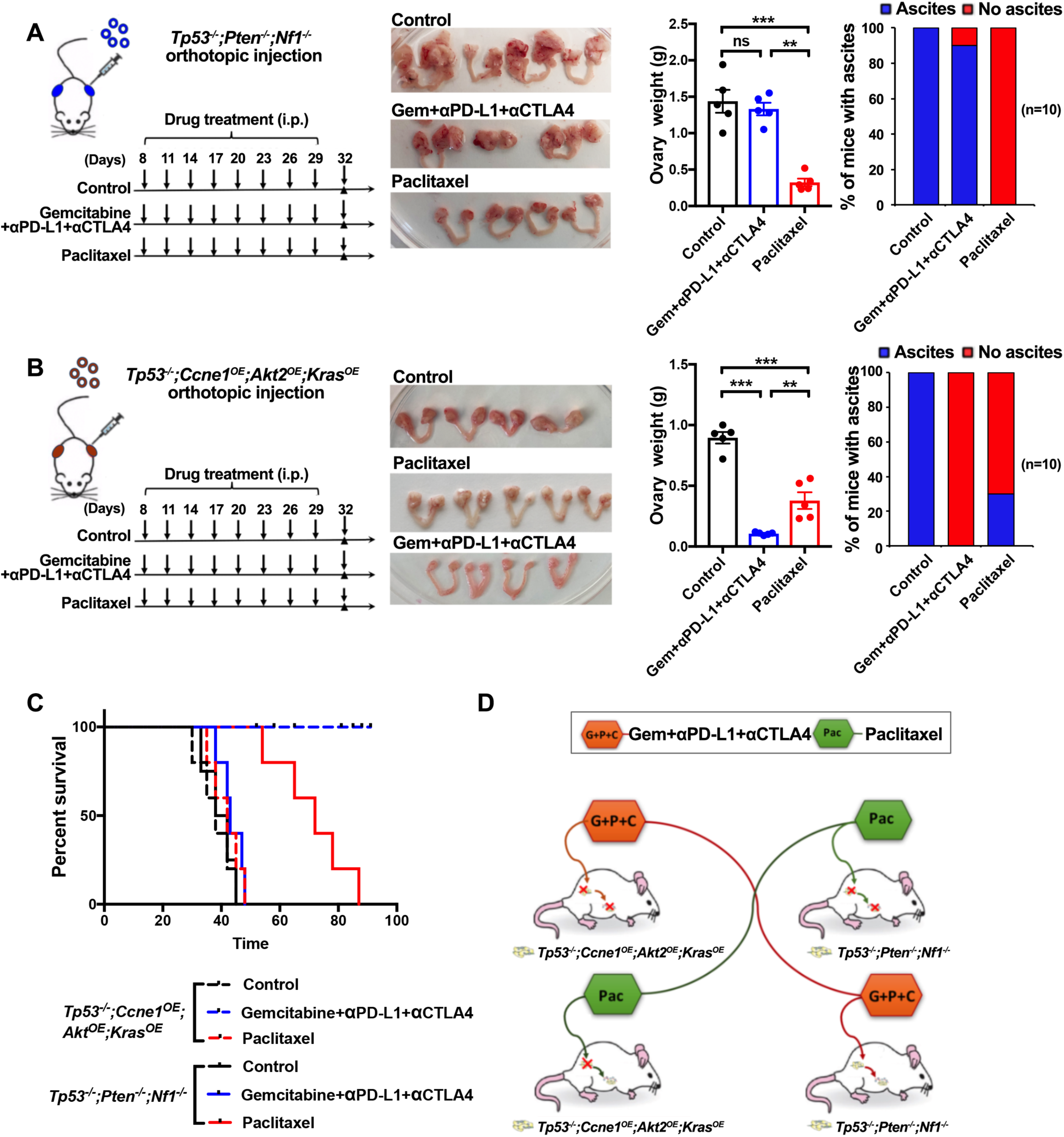
Treatment efficacy is tumor genotype-dependent. (A) Left panel: Schematic showing treatment of *Tp53*^*-/-*^;*Pten*^*-/-*^;*Nf1*^*-/-*^ tumor-bearing mice with Gemcitabine/α-PD-L1/α-CTLA-4 regimen (from Figure 6) or Paclitaxel. Second panel: Genital tracts from mice treated as indicated; Third panel: Ovary weights in treated mice. Right panel: % mice with ascites after indicated treatment. (B) Left panel: Schematic showing treatment of *Tp53*^*-/-*^;*Ccne1*^*OE*^;*Akt2*^*OE*^;*Kras*^*OE*^ tumor-bearing mice with the indicated regimens. Second panel: Genital tracts from mice treated as indicated; Third panel: Ovary weights in treated mice. Right panel: % mice with ascites after indicated treatment. Data indicate means ± SEM, **p < 0.01, unpaired t test. (C) K-M curves of tumor-bearing *Tp53*^*-/-*^;*Pten*^*-/-*^;*Nf1*^*-/-*^ or *Tp53*^*-/-*^;*Ccne1*^*OE*^;*Akt2*^*OE*^;*Kras*^*OE*^ mice, treated as indicated. Treatments were withdrawn at Day 32. (D) Cartoon summarizing results, depicting tumor genotype-specificity of therapeutic efficacy. See also Figure S7.

Both showed some response, but as predicted by our *in vitro* experiments, mice bearing *Tp53*^*-/-*^;*Pten*^*-/-*^;*Nf1*^*-/-*^ tumors had a greater therapeutic response and survived much longer than those with *Tp53*^*-/-*^;*Ccne1*^*OE*^;*Akt2*^*OE*^;*Kras*^*OE*^ tumors (Figure 7B and 7C). Although single agent paclitaxel did not result in CRs, it evoked several potentially beneficial changes in the TME, including an influx of CD4 and CD8 T cells with signs of activation (CD44+) and lower levels of exhaustion markers, as well as a decrease in g-MDSC and macrophages, with those remaining showing increased M1-like phenotype (Figures S7A-C).

## DISCUSSION

Like most solid tumors, HGSOC is a genetically complex, diverse disease, yet with the exception of PARP-Is for *BRCA*-mutant tumors, current therapy for HGSOC (and for most other neoplasms) remains genotype-agnostic. Perhaps unsurprisingly, ∼20% of HGSOC patients experience minimal or no clinical benefit from such uniform treatment; nearly all of the rest relapse and die after an initially positive response (Binnewies et al., 2018). Rational development of genotype-informed therapies for HGSOC is impeded by a paucity of relevant experimental models that allow conventional, targeted, and immune therapies to be tested alone or in combination. Current immune-competent mouse models fail to represent the genetic diversity of HGSOC, and the most often-used syngeneic model (ID8) initiates from the less common cell-of-origin (OSE) and lacks the genetic abnormalities seen in the disease. Our FTE organoid-based system remedies these deficiencies, enabling analysis of the consequences of specific genetic aberrations on *in vitro* properties (proliferation, differentiation, morphology, drug sensitivity, secretome), assignment of complementation groups for tumorigenicity and metastasis, assessment of the TME, and evaluation of drug therapies (Supplementary Figure 1). In contrast to the “field effect” seen in conventional GEMMs, organoid-derived tumors form in the correct anatomical location, surrounded by normal host cells. We demonstrate the utility of this platform by developing a combination regimen that evokes durable complete responses in mice bearing CCNE1-overexpressing tumors, the human counterparts of which are refractory to standard therapy, but which has no activity against *Tp53*^*-/-*^;*Pten*^*-/-*^;*Nf1*^*-/-*^ tumors. By contrast, the latter are unexpectedly more sensitive to paclitaxel (Figure 7D). These data argue strongly against therapeutic approaches that treat HGSOC as a single entity. While our focus was on HGSOC originating from FTE, our platform can be extended easily to OSE-derived HGSOC, as well as to any of the increasing number of tumor types for which mouse organoids can be propagated and engineered (Clevers, 2016; Drost and Clevers, 2018; Neal and Kuo, 2016).

The cell-of-origin for HGSOC remains controversial. Although transcriptomic (Ducie et al., 2017; Hao et al., 2017; Lawrenson et al., 2019), proteomic (Coscia et al., 2016), epigenomic (Lawrenson et al., 2019), and mouse modeling studies (Szabova et al., 2012; Zhang et al., 2019) suggest that at least some cases initiate in OSE, most HGSOC probably initiates in FTE (Ducie et al., 2017; Karnezis and Cho, 2017; Karnezis et al., 2017). For this reason, we engineered our initial models using FTE organoids. Nevertheless, orthotopic injection of 10^5^ engineered *Tp53*^*-/-*^;*Brca1*^*-/-*^;*Myc*^*OE*^ OSE cells also gives rise to HGSOC-like tumors, which kill recipients within 50 days (Ding et al., 2018). Notably, mice injected with more FTE cells (2×10^6^) bearing the same mutations survive from 70 days-150 days, consistent with our previous finding that the cell-of-origin of HGSOC influences tumor biology (Zhang et al., 2019).

Tumorigenic organoids showed several expected, but other unanticipated, sensitivities to small molecule inhibitors/drugs. In line with previous studies of conventional ovarian cancer cell lines, and *Tp53*^*-/-*^;*Brca1*^*-/-*^;*Myc*^*OE*^ OSE-derived cells, *Brca1-*mutant FTE-derived tumor organoids have increased PARP-I sensitivity. Notably, the degree of hypersensitivity tends to be greater in conventional *Brca*-mutant cell lines (Pulliam et al., 2018), than in our FTE-derived tumorigenic organoids. Conceivably, high MYC levels modulate PARP-I response; indeed, it will be important to test the effects of other genetic abnormalities that commonly co-occur with *Brca1* mutations on PARP-I sensitivity. Moreover, while there was a class-specific increase in PARP-I sensitivity in *Tp53*^*-/-*^;*Brca1*^*-/-*^;*Myc*^*OE*^ organoids, the extent of hypersensitivity differed for individual PARP-Is. Future studies will aim to elicit the mechanistic basis for these differences, as well as their respective effects on the TME. ATR inhibitors also showed increased efficacy against *Tp53*^*-*^ */-;Brca1*^*-/-*^;*Myc*^*OE*^ organoids, in accord with the HR-deficiency conferred by BRCA1 deficiency. Likewise, the increased sensitivity of *Tp53*^*-/-*^;*Pten*^*-/-*^;*Nf1*^*-/-*^ cells to ATR inhibition comports with the previously reported role for nuclear PTEN in HR (Bassi et al., 2013; Shen et al., 2007). The reasons for the genotype-dependent differences in sensitivity to paclitaxel (for *Tp53*^*-/-*^;*Pten*^*-/-*^;*Nf1*^*-/-*^ cells) and to chloroquine (for *Tp53*^*-/-*^;*Pten*^*-/-*^;*Nf1*^*-/-*^>*Tp53*^*-/-*^;*Ccne1*^*OE*^;*Akt2*^*OE*^;*Kras*) are less clear. Comparison of *Tp53*^*-/-*^;*Pten*^*-/-*^ and *Tp53*^*/--*^;*Nf1*^*-/-*^ organoids implicate NF1 deficiency as the main cause increased paclitaxel sensitivity (data not shown); notably, several studies indicate that NF1 associates with microtubules (Gregory et al., 1993; Roby et al., 2000; Xu and Gutmann, 1997), the target of paclitaxel. Previous studies demonstrated that *KRAS*-mutant cells require autophagy for survival (Rosenfeldt et al., 2013), and PTEN deficiency or AKT2 over-expression, by increasing mTOR activation, would be expected to suppress autophagy. Conceivably, increased RAS activation caused by NF1 deficiency or KRAS over-expression, combined with basal autophagy suppression due to increased mTOR activity, sensitizes FTE organoids to further inhibition of autophagy by chloroquine. Whatever the exact mechanisms, these differences in drug sensitivity emphasize the value of more precise, genotype-informed models for developing new therapies. Although we only tested a small subset of agents, our models can easily be used for high throughput drug screens or genetic perturbations (e.g., CRISPR/Cas9 screens). Furthermore, the genotype-specific sensitivities that we observe suggest that only certain subsets of patients will respond to specific standard-of-care single agent or combination therapies. For example, most of the benefit of adding paclitaxel to platinum, a practice that developed empirically (Boyd and Muggia, 2018; Kampan et al., 2015), might accrue mainly to those patients with PTEN-deficient tumors; other patients might only incur taxane-based toxicity.

Studies of human ovarian cancer also reveal a complex TME, with differences in infiltrating immune cells and tumor-associated chemokines/cytokines variably associated with prognosis (Rodriguez et al., 2018). For example, as in many other malignancies, intra-tumor CD8+ and high CD8+/Treg cells are associated with improved survival, whereas high levels of Tregs are a negative prognostic sign (Curiel et al., 2004; Hwang et al., 2012; Stumpf et al., 2009; Zhang et al., 2003). In turn, intra-tumor T cells have been associated with expression of *CXCL9, CXCL10, CCL5, CCL21*, and/or *CCL22*, whereas high *VEGF* levels inversely correlate with T cell infiltration (Bronger et al., 2016; Nagarsheth et al., 2017; Zhang et al., 2003). A large pan-cancer genomic analysis indicated that high levels of CCL5 RNA and protein (by IHC) correlate with intra-tumor CD8 cells in solid tumors, including ∼75% of ovarian carcinomas (Negus et al., 1997). *CCL5* and *CXCL9* also correlated in this analysis, and dual expression of these chemokines was associated with better prognosis. Interestingly, ovarian cancers with high intra-tumor CD8+ cells but low *CCL5* had high levels of *CXCL9* transcripts (Dangaj et al., 2019). By contrast, high levels of tumor-associated macrophages, particularly M2-like macrophages, and MDSCs correlate with poor outcome (Drakes and Stiff, 2018; Ruffell and Coussens, 2015). Aside from describing greater T cell infiltration and better prognosis in *BRCA*-mutant tumors (Clarke et al., 2009; Rodriguez et al., 2018; Strickland et al., 2016) and examining the regulatory mechanism of specific immune regulatory molecules (e.g., silencing of CCL5 transcription) (Dangaj et al., 2019), these studies are tumor genotype-agnostic. Yet the three mouse models that we examine in detail display major differences in TME, as well as significant differences in chemokine and cytokine gene expression, including tumors where CXCL10 likely directs T cell influx, and others in which T cell influx is associated with *Cxcl9*. As tumor genotype will affect response to targeted and conventional agents, understanding how genotypic differences direct host immune responses could aid in therapy development. Our ability to manipulate tumor (e.g., by further engineering of chemokine/cytokine genes) and host immune cells (e.g., by depletion studies, injection of tumorigenic organoid cells into various knockout backgrounds), and to study tumors over time, should provide insights into how the ultimate TME develops and respond to therapy.

Previous studies have also reported model-specific differences in tumor immune infiltrates (Jiao et al., 2019; Li and Stanger, 2019). For example, MYC, KRAS, mTOR, Yes-associated protein (YAP), and β-catenin signaling in cancer cells all have been reported to influence the TME (Wellenstein and de Visser, 2018). Many of these studies examined specific GEMMs, syngeneic tumor models, or GEMM-derived cell lines, some engineered to have additional genetic abnormalities not seen in the parental model. Potentially causal cytokines/chemokines that alter the TME have also been proposed or identified in such studies. However, the extent to which TME responses are “hard-wired” by specific oncogenic defects in tumors is unclear. For example, PTEN deficiency leads to impaired T cell infiltration owing to immunosuppressive myeloid cells in mouse models of prostate cancer (Wang et al., 2016) and melanoma (Peng et al., 2016). But whereas CXCL5 (mouse)/CXCL6 (human) are implicated in myeloid cell immigration in prostate cancer, increased CCL2 and VEGF appear to be the culprits in melanoma. We also observed increased myeloid cells in *Tp53*^*-/-*^;*Pten*^*-/-*^;*Nf1*^*-/-*^ HGSOC, along with increased levels of CCL2 and VEGF. However, MCSF1 and CXCL1 might also contribute to myeloid infiltration in this model, whereas CXCL5 is not elevated and is unlikely to play a role. These findings, along with many others (Li and Stanger, 2019; Spranger and Gajewski, 2018; Wellenstein and de Visser, 2018) argue that cellular context (e.g., cell-of-origin, cooperating mutations) might be equally important as any one specific oncogenic abnormality for determining the ultimate TME and anti-tumor immune response. Our ability to rapidly engineer FTE and OSE organoids with all major combinations of genetic defects seen in HGSOC will allow us to address this important issue.

Attempts to manage HGSOC with immune therapy have not been very fruitful. Single-agent anti-CTLA-4 or anti-PD1/PD-L1 yield only modest results, with response rates of 10%-15% (Hamanishi et al., 2015; Liu et al., 2019). Combining anti-PD1 and anti-CTLA4 increases response rate to 34%, but the clinical data are very immature (Ghisoni et al., 2019). *BRCA1*-mutant HGSOC shows greater T cell infiltration, as does our *Brca1*-mutant mouse model, and such tumors might show a better response to immune checkpoint inhibition, alone or in combination with PARP-Is (Ding et al., 2018; Shen et al., 2019; Strickland et al., 2016). However, there is no evidence that these responses are durable, and whether other tumor genotypes confer more or less sensitivity is not clear. A major advantage of our organoid platform is its ability to rapidly suggest and test potential therapies. Our chemo-immunotherapy combination of three approved drugs, gemcitabine, anti-PD-L1, and anti-CTLA4, leads to durable, T-cell-dependent CRs in a highly aggressive, CCNE1-overexpressing model of HGSOC. Importantly, patients with such tumors are highly refractory to current therapies (Au-Yeung et al., 2017; Nakayama et al., 2010). It will be important to test the generality of these responses in our other models in which *Ccne1* over-expression is accompanied by other genetic defects (e.g., *Mecom* and/or *Myc* over-expression), as well as to develop and test combination immunotherapies with paclitaxel in PTEN/NF1-deficient HGSOC. Indeed, the changes in the TME that ensue following single agent treatment suggest several drugs to combine with paclitaxel in this model. In conclusion, our ability to rapidly generate multiple, genetically defined, complex HGSOC organoid models should facilitate studies of the diversity and host response of this disease. Our models also suggest a genomics-informed, rationally based combination treatment for *CCNE1*-amplified HGSOC, and suggest new interventional strategies for other genomic subgroups of this highly complex disease.

## STAR⋆Methods

### Key Resources Table

See attached Excel sheet

### Lead Contact and Materials Availability

Further information and requests for resources and reagents should be directed to, and will be fulfilled by, the Lead Contact, Dr. Benjamin G. Neel.

### Experimental Model and Subject Details

*Tp53*^*f/f*^ (BALB/c) female mice were a gift from Dr. Kwok-kin Wong (NYUSoM). *Tp53*^*f/f*^;*Brca1*^*f/f*^ mice (B6) (Liu et al., 2007) were provided by Dr. Richard Possemato (NYUSoM). Female C57BL/6 mice (6-8 weeks old) were purchased from Charles River Laboratories. All animal experiments were approved by, and conducted in accordance with the procedures of, the IACUC at New York University School of Medicine (Protocol no.170602).

Where indicated, mice were injected intraperitoneally (i.p.) with gemcitabine (50 mg/kg), paclitaxel (40 mg/kg), anti-CTLA4 (50 µg, clone 9H10, BioXcel), and/or anti-PD-L1 (50 µg, clone 4H2, BioXcel), beginning 8 days after tumor cell implantation. Dosing was repeated every three days for the duration of the experiment. Control mice were injected with PBS or isotype control antibody (clone LTF-2, BioXcel).

CD4^+^ and/or CD8^+^ T cells were depleted by i.p. injection of 200 µg of *InVivo*MAb anti-mouse CD4 (clone GK1.5, BioXcel) and/or *InVivo*MAb anti-mouse CD8α (clone 2.43, BioXcel) respectively, one week after tumor implantation. Injections were repeated every 3 days for the duration of the experiment (Li et al., 2018). Other mice were injected with isotype control antibody (clone LTF-2, BioXcel). Depletion of the appropriate lymphoid population was confirmed by flow cytometric analysis of the peripheral blood and reassessed every 2 weeks for the duration of the study. Peripheral blood was collected from tail veins into heparinized microhematocrit capillary tubes, separated by centrifugation, and prepared for flow cytometry by treatment with ACK lysis buffer to remove red blood cells, followed by serial washes in RPMI.

## Detailed Methods

### Organoid culture and engineering

Fallopian tubes from *Tp53*^*f/f*^ or *Tp53*^*f/f*^;*Brca1*^*f/f*^ mice were collected, freed from surrounding tissue under a dissecting microscope (Leica), minced, digested with collagenase type I and Dispase 0.012% (w/v) at 37°C for 1h, and incubated with TryLE™ express (Thermo Fisher Scientific) for 10 mins at 37°C. Cells were passed through a strainer (70 µm) and seeded into EGF-reduced Matrigel (BD Bioscience) in fallopian tube organoid medium, as described (Zhang et al., 2019). All cultures were checked for mycoplasma monthly using a PCR assay. Organoid cells were collected by using cold Cultrex® Organoid Harvesting Solution (Stem Cell Technologies) to dissolve Matrigel, following the manufacturer’s instructions.

For *in vitro* deletion of *Tp53* and/or *Brca1, Tp53*^*f/f*^ female or *Tp53*^*f/f*^;*Brca1*^*f/f*^ organoids were dissociated into single cells as described (Koo et al., 2011), and then infected with 10^5^ pfu Adenovirus-CMV-Cre (Vector Development Lab, Baylor College of Medicine) by “spinoculation” at 37 for 1h. Cell pellets were collected and seeded into Matrigel, and 7 days later, organoids were released as described above, and single organoids were picked and expanded. Deleted clones were identified by PCR. Primers were described previously (Liu et al., 2007); their sequences are provided in the Supplementary Information. MYC overexpression was detected by immunoblotting.

Mouse *Ccne1* and *Akt2* were cloned into pLJM1 (Addgene: #19319), modified with neomycin-resistance or blasticidin-resistance genes, respectively. For *Myc*-over-expression, we used the vector MSCV-transgene-PGK-Puro IRES-GFP, purchased from Addgene (#75124). Mouse *Kras* was cloned into pMSCV-IRES-mCherry (Addgene, #52114). Successful gene insertion was confirmed by Sanger sequencing. *Pten* (agatcgttagcagaaacaaa) or *Nf1* (ctcgtcgaagcggctgacca) sgRNA sequences were designed with the CRISPR Design Tool (http://crispr.mit.edu/) and inserted into LentiCRISPR v2 (Addgene, #52961). For virus production, lentiviral vectors were co-transfected with psPAX2 and pMD2.G into HEK293T cells at a ratio of 10:7.5:2.5 or retroviral vectors were co-transfected with pVPack and VSV-G into HEK293T cells at a ratio of 10:6.5:3.5. All transfections were performed by using Lipofectamine 2000 Transfection Reagent (Thermo Fisher Scientific), according to the manufacturer’s instruction. Media were changed 8h after transfection, and viral supernatants were collected 48h later by passage through a 0.45-mm filter, aliquoted, and stored at −80°C.

Organoids were dissociated into single cells and spinoculated with lentiviruses/retroviruses, also as described (Koo et al., 2011). Briefly, viral supernatants were added to cells in 48-well plates, centrifuged at 600g at 37°C for 60 min, incubated at 37°C for another 6-8 hours, collected, and re-seeded in Matrigel-containing media. Infected organoids were selected 72 hours after viral transduction with G418 (Thermo Fisher, 10131027) or blasticidin (Sigma, 15205), as indicated. Gene deletion and/or overexpression was assessed by PCR or immunoblotting. At least two independent clones of each genotype were used for experiments.

For orthotopic tumorigenicity assays, cell pellets were collected and injected into ovarian bursae, as described (Zhang et al., 2019). Briefly, mice were anesthetized by intraperitoneal injection of Xylazine (10 mg/kg) and Ketamine (50 mg/kg), shaved, and cleaned with betadine. A dorsal incision above the ovary was made, followed by incision of the peritoneal cavity. The ovary was externalized, and cells (2 × 10^6^) in 50 µl PBS/Matrigel (1:1 v/v) were injected through the ovarian fat pad to the bursa using an insulin syringe with a 31G needle. Injected ovaries were returned to the peritoneal cavity, and incisions were sealed with wound clips. Mice that developed tumors were euthanized at the indicated time(s), or for survival experiments, they were monitored until death or upon veterinary recommendation.

### Flow Cytometry

Tumors were minced, chopped, and digested with Gentle Collagenase, Dispase 0.012% (w/v) and DNaseI (STEMCELL Technologies) at 37°C for 1 hour. Single cell suspensions were obtained by passage through a strainer (70 µm), washed in FACS buffer (PBS with 5% FBS), incubated with LIVE/DEAD Fixable Zombie Yellow Fixable Viability Kit (Biolegend, 423104) for 30 min., and then blocked with anti-CD16/32 (Biolegend, clone 93) for 5 min. on ice. Samples were then incubated with primary fluorophore-conjugated antibodies on ice for 45 min. For detection of intracellular markers, FOXP3 Fixation/Permeabilization Buffer Set (BioLegend) was used, according to the manufacturer’s instructions. Antibodies for flow cytometry are described in the Star Methods. Flow cytometry was performed on an LSR II flow cytometer at the Flow Cytometry Core of the PCC Precision Immunology Shared Resource and analyzed with FlowJo software. Organoids cultured 6 days after infection with MSCV-Kras-mCherry were collected and digested as above, passed through a strainer (70 µm) to obtain single-cell suspensions, centrifuged at 1000×g for 5 min, and resuspended in PBS containing 2% FBS, 10 µM Y-27632, (STEMCELL Technologies Inc.), and DAPI (1 µg/ml). FACS was performed immediately on a MoFloTM XDP, and mCherry^hi^ and mCherry^neg^ cells were seeded at 5,000/well.

### Bioluminescence Imaging

Mice were injected with 150 mg/kg D-luciferin Firefly (PerkinElmer Part Number #122799), and luminescence was assessed 15 min. later by using a PerkinElmer IVIS Lumina III imaging system. Images were analyzed with Living Image Software 4.7.3.

### Cytokine Profiling

Cytokine levels from conditioned media of cultured tumorigenic organoid cells (72h) were profiled using services at Eve Technologies (Calgary, Canada). The Mouse Cytokine Array/Chemokine Array 31-Plex (MD31) was used at the recommended dilutions; this panel includes Eotaxin (CCL11), G-CSF, GM-CSF, IFNγ, IL-1α, IL-1β, IL-2, IL-3, IL-4, IL-5, IL-6, IL-7, IL-9, IL-10, IL-12 (p40), IL-12 (p70), IL-13, IL-15, IL-17A, IP-10, KC (CXCL1), LIF, LIX (CXCL5), MCP-1 (CCL2), M-CSF, MIG (CXCL9), MIP-1α (CCL3), MIP-1β (CCL4), MIP-2 (CXCL2), RANTES (CCL5), TNFα, and VEGF.

### Drug sensitivity assays

Tumorigenic organoid cells were seeded into 96-well plates at 1,000 cells/well (Day 0). The indicated concentrations of Rucaparib, Niraparib, Olaparib, Gemcitabine, Doxorubin, Paclitaxel, Carboplatin, Seliciclib, PHA767491, BAY1895344, Chloroquine and YKL-5-124 were added on the day following seeding (Day 1). Media were changed, and fresh drug was added on Day 3. Cell viability was assessed on Day 5 by adding10 μl PrestoBlue and incubating for 30 min in 37°C. Fluorescence was measured in a FlexStation® 3 Multi-Mode Microplate Reader (BOSTONind). Results were normalized to DMSO controls, and IC_50_ values were determined using Graphpad Prism 7.

### Histology and imaging

Tumor tissues were fixed in 4% PFA for 48 hours, followed by processing and embedding for standard histological analysis, immunohistochemistry (IHC) or immunofluorescence (IF). Sections (5 µm) were de-paraffinized, rehydrated, and stained with hematoxylin and eosin (H&E) or were subjected to antigen retrieval (citrate) at 120°C in a pressure cooker for 15 minutes for 5 minutes. For IHC, endogenous peroxidase activity was quenched in 3% H_2_O_2_ in methanol for 15 min, and sections were blocked in 0.5% BSA in PBS for 1h. Primary antibodies were added overnight at 4°C. Slides were then washed in PBS three times for 10 min, incubated with secondary antibodies for 1 h at room temperature, and washed again. Antigens were detected by using the HRP Polymer Detection Kit and DAB peroxidase (HRP) substrate (34002, Life Technologies). For IF, after antigen retrieval, slides were washed in PBS three times for 10 min and then blocked in 0.5% BSA in PBS for 1h. Primary antibodies were incubated at 4 °C overnight, and sections were washed in PBS (3 times, 10 min each), followed by incubation with Alexa Fluor secondary antibodies, as indicated. After washing, slides were mounted with Prolong™ Gold Antifade Mountant (Thermo Fisher, P36930) for imaging. For IF staining, organoids were released from Matrigel (as above), transferred to a µ-Slide 8 Well Glass Bottom (Ibidi), fixed in 4% paraformaldehyde (pH 7.4) for 20 min, permeabilized in 1% Triton X-100 in PBS, and blocked with PBS, 1% BSA, 3% normal goat serum, 0.2% Triton X-100. Following incubation with primary antibody at 4°C overnight, organoids were washed in PBS three times for 10 min and incubated at room temperature with the appropriate Alexa Fluor secondary antibody (1:200). Organoids were washed with PBS and mounted using ProLong gold antifade (Molecular Probes, Invitrogen). IHC slides were scanned by using a Leica SCN400 F whole-slide scanner. IF images were taken with a confocal microscope (ZEISS LSM 700).

### RNA Extraction and Sequencing

Tumors were collected in Trizol, and RNA was extracted using the miRNeasy Mini Kit (Qiagen), according to the manufacturer’s instructions. RNA sequencing was performed by the PCC Genome Technology Center Shared Resource (GTC). Libraries were prepared by using the Illumina TruSeq Stranded Total RNA Sample Preparation Kit and sequenced on an Illumina NovaSeq 6000 using 150 bp paired-end reads. Sequencing results were de-multiplexed and converted to FASTQ format using Illumina bcl2fastq software. The average number of read pairs/sample was 35.4 million. Data were processed by the Perlmutter Cancer Center Applied Bioinformatics Laboratories shared resource (ABL). Briefly, reads were adapter- and quality-trimmed with Trimmomatic (Bolger et al., 2014) and then aligned to the mouse genome (build mm10/GRCm38) using the splice-aware STAR aligner (Dobin et al., 2013). The featureCounts program (Liao et al., 2014) was utilized to generate counts for each gene, based on how many aligned reads overlap its exons. These counts were normalized and tested for differential expression, using negative binomial generalized linear models implemented by the DESeq2 R package (Love et al., 2014). For pairwise differential expression analysis between the tumor groups, normal FT samples were not included in the model. Statistical analysis and visualization of gene sets were performed using the fgsea (Korotkevich G., 2019) and clusterProfiler R packages (Yu et al., 2012).

### Quantification and Statistical Analysis

Unless otherwise specified, data are presented as means ± SEM. Survival rates were analyzed by log-rank test, using GraphPad Prism software. Unless otherwise specified, P values were determined by two-tailed Student’s t-test, with P< 0.05 considered statistically significant.

### Data availability

RNA sequence data have been deposited in the GEO database under the accession code GSE147276. All other data supporting the findings of this study are available within the article, the Supplemental information files, or the corresponding author upon request.

## Acknowledgements

We thank the PCC Experimental Pathology, Precision Immunology, Microscopy, GTC, Preclinical Imaging Laboratory, and ABL shared resources for technical support, and Drs. Justin Mastroianni (PCC) for assistance with IVIS imaging. We thank Drs. Kwan Ho Tang, Mitchell Geer, and Carmine Fedele (Neel lab) for helpful advice and discussion. Initial work on this project was supported by grant-MOP-191992 from the Canadian Institutes for Health Research to B.G.N. S.Z. was supported by a post-doctoral fellowship from Ovarian Cancer Research Fund Alliance. PCC shared resources are supported by P30 CA01687.

## Author Contributions

S.Z., S.I., B.G.N., and R.A.W. designed the study. S.Z. performed the majority of the experiments. S.I. provided some plasmid constructs and virus preparations. I.D. performed the bioinformatics analyses. H.R. assisted with orthotopic injections and *in vivo* drug delivery. W.W performed the dose-response experiments. C.F. helped with T cell depletion experiments. S.Z., I.D., and B.G.N., wrote the manuscript, which was edited by S.Z., I.D., B.G.N., S.I., and R.A.W.

## Supplemental Methods

### Primer sequences used for genotyping

*Brca1 fl:* Genotyping floxed forward: 5’-TATCACCACTGAATCTCTACCG -3’; Genotyping floxed reverse: 5’-GACCTCAAACTCTGAGATCCAC -3’; floxed alleles: 545 bp, wild type alleles: 390 bp

*Brca1Δ:* Genotyping floxed forward: 5’-TATCACCACTGAATCTCTACCG -3’; Genotyping floxed reverse: 5’-TCCATAGCATCTCCTTCTAAA C -3’. Deleted fragment: 594 bp.

## Supplemental Figures

**Figure S1.**
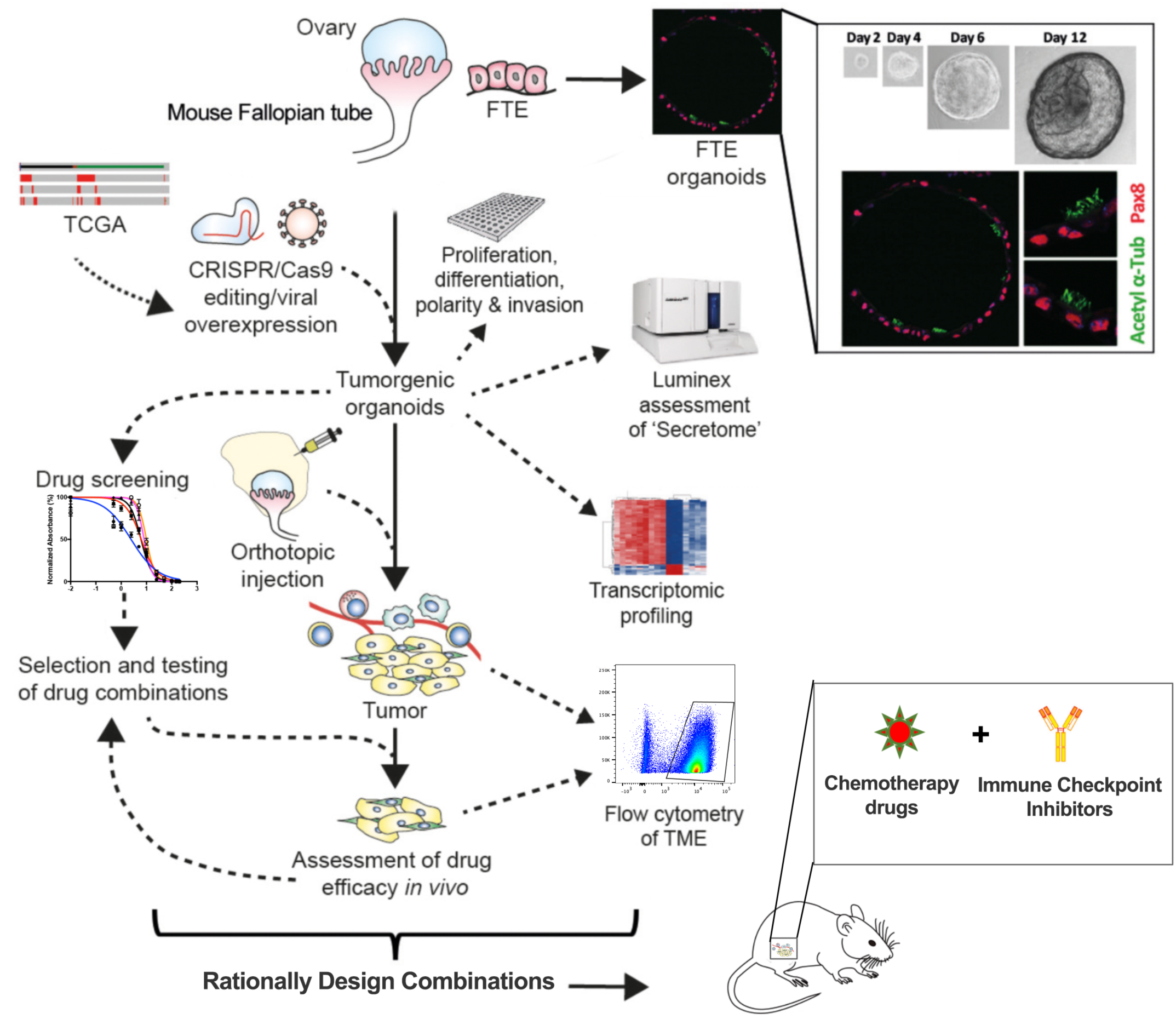
Schematic depicting organoid platform, related to Figure 1.

**Figure S2.**
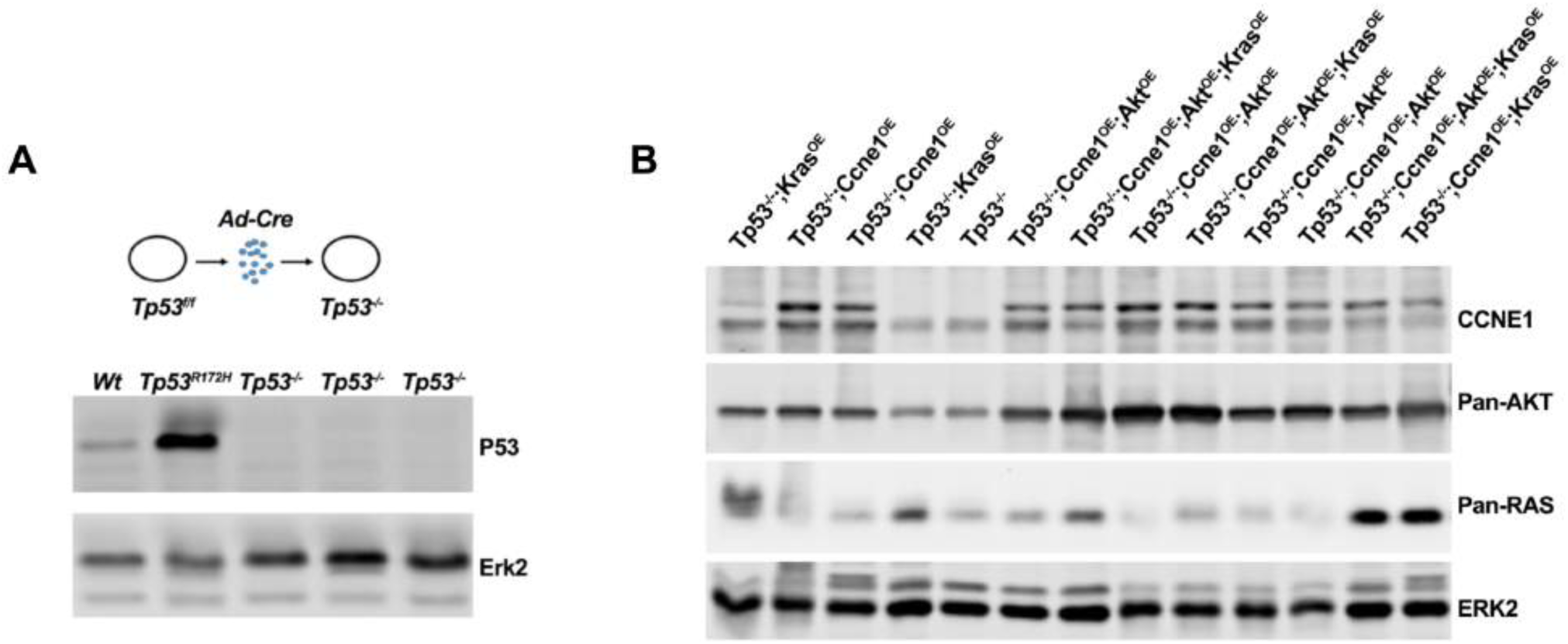
Confirmation of successfully engineered organoids, related to Figure 2 and 3. (A) Immunoblot of organoid lysates showing effective deletion of floxed *Tp53* alleles following adeno-Cre infection (*Tp53*^*-/-*^). Lysates from wild type (WT) and *Tp53*^*R172H/+*^ organoids are shown as controls. (B) Immunoblot showing expression of the indicated proteins in engineered organoids. Genotypes are shown at top; each lane represents a different clone. ERK2 serves as a loading control.

**Figure S3.**
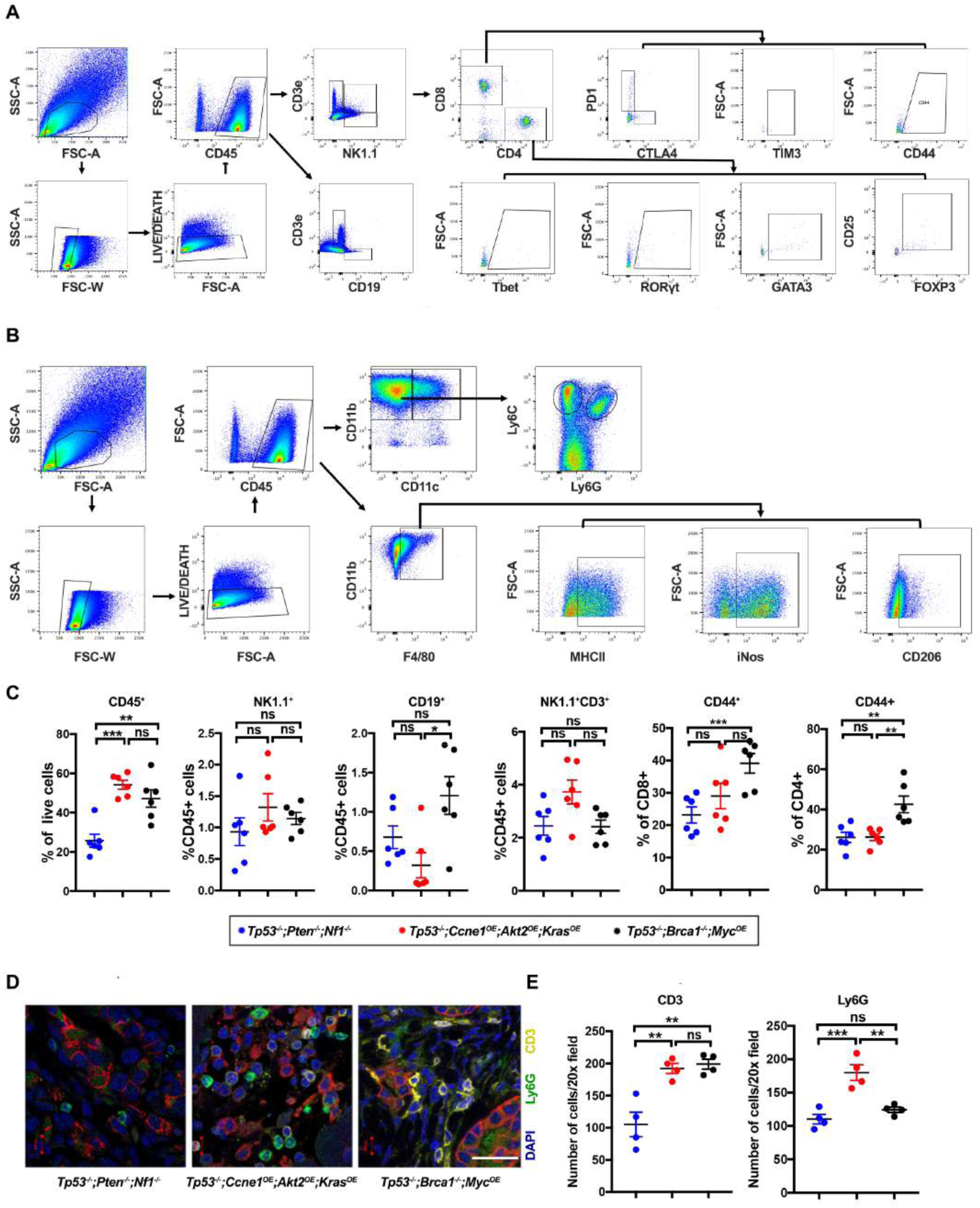
Immune landscape analysis, related to Figure 4. (A) Gating strategy for lymphoid cells. (B) Gating strategy for myeloid cells. (C) Analysis of tumors of different genotypes with the indicated markers/marker combinations. Each point represents a tumor from an independent mouse. Horizontal bar indicates mean cell percentage; error bars indicate ± SEM; **P<0.01, ***P<0.001, 2-way ANOVA. (D) Representative immunofluorescence micrograph showing staining for CD3 (yellow), Ly6G (green), Cytokeratin 7 (red), and nuclei (blue) in tumors of the indicated genotype. (E) Quantification of CD3+ and Ly6G+ cells in tumors of the indicated genotypes, averaged over five 20X fields. Horizontal lines in each panel indicate the mean cell number; error bars represent ± SEM, **P<0.01, ***P<0.001, 2-way ANOVA.

**Figure S4.**
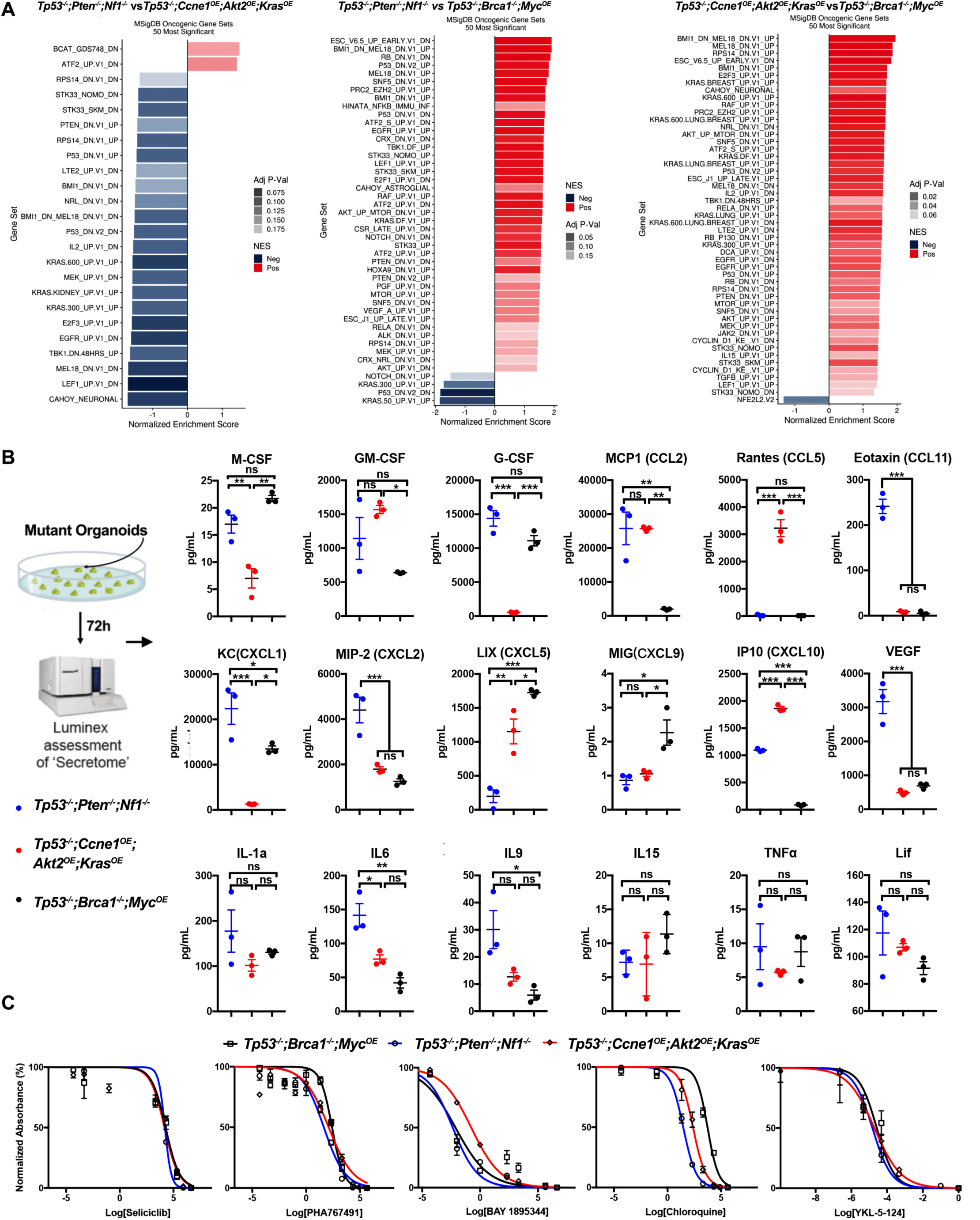
Further characterization of tumorigenic organoids and tumors, related to Figure 4 and 5. (A) Pathway analysis comparing the indicated groups. Significantly enriched KEGG (left) and MSigDB Hallmark genes (right) of the genes ranked by fold-change between the indicated groups are shown. Shading represents the FDR-adjusted p-value within each category; color indicates direction of enrichment relative to the first group of the comparison. (B) Level of expression of indicated cytokines, chemokines and growth factors. error bars indicate ± SEM; **P<0.01, ***P<0.001, 2-way ANOVA. (C) Dose-response curves for tumorigenic organoids treated with the indicated targeted agents.

**Figure S5.**
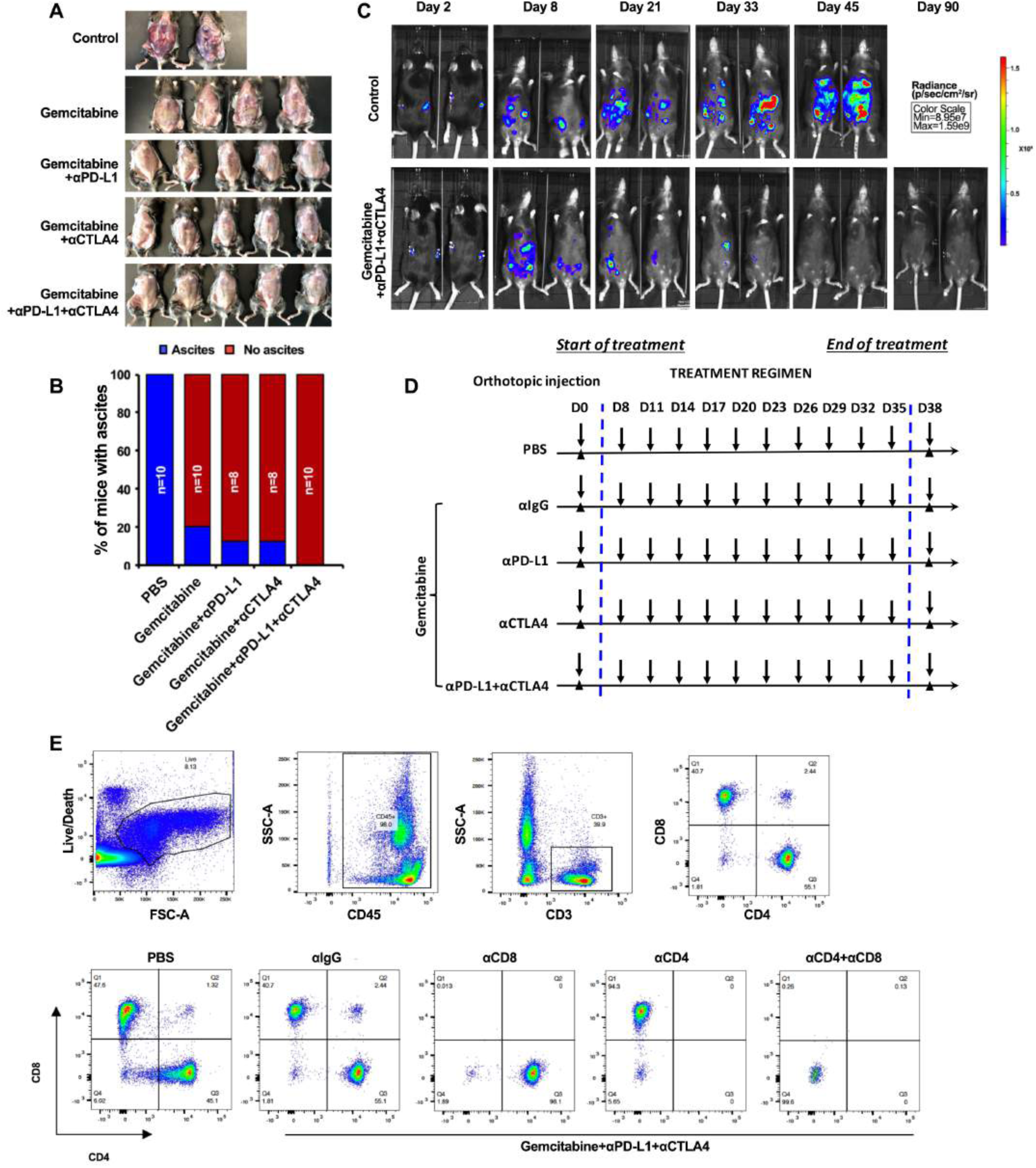
Response of *Tp53*^*-/-*^;*Ccne1*^*OE*^;*Akt2*^*OE*^;*Kras*^*OE*^ tumor-bearing mice to combination therapy is T cell-dependent, related to Figure 6. (A) Exposed abdominal cavities of mice treated as indicated. Note distention due to ascites in all but the 3-drug combination-treated mice, and large ovarian tumor in the PBS group, as indicated in Figure 6A. (B) Percentage of mice with ascites following the indicated treatment regimens, assessed at Day 35; see Fig. 6A. (C) Representative bioluminescence images of mice bearing orthotopic tumor allografts (luciferized) with indicated treatments measured at day 2, 8, 21, 33, 45 and 90, respectively. Note that treatment was terminated after day 35, and there was no recurrence over the monitoring period (90 days). (D) Schematic depicting treatment regimens, n=5 mice/group. (E) Representative flow cytometric analysis of peripheral blood leukocytes, demonstrating depletion of the indicated T cell populations.

**Figure S6.**
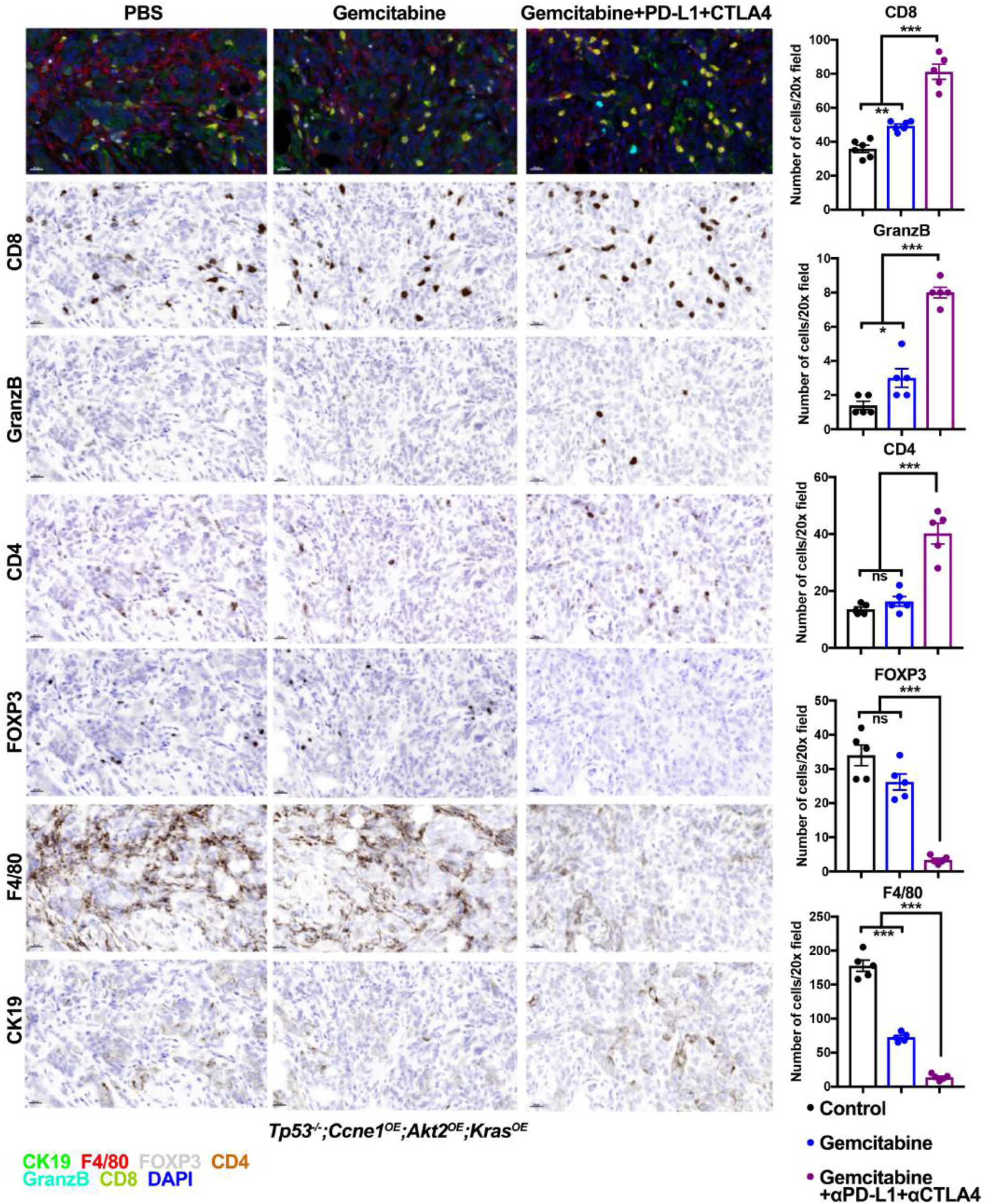
Multiplex IHC analysis of *Tp53*^*-/-*^;*Ccne1*^*OE*^;*Akt2*^*OE*^;*Kras*^*OE*^ tumors treated with PBS, Gemcitabine, or Gemcitabin+aCTLA4+aPD-L1, related to Figure 6. Left panels: sections were stained with the indicated antibodies and analyzed by OPAL. scale bars: 20 µm. Right panels: quantification of in indicated tumors. Quantification of the indicated markers in tumors of the indicated genotype from average cell numbers from five 20X fields. Data represent mean ± SEM, **P<0.01, ***P<0.001, 2-way ANOVA.

**Figure S7.**
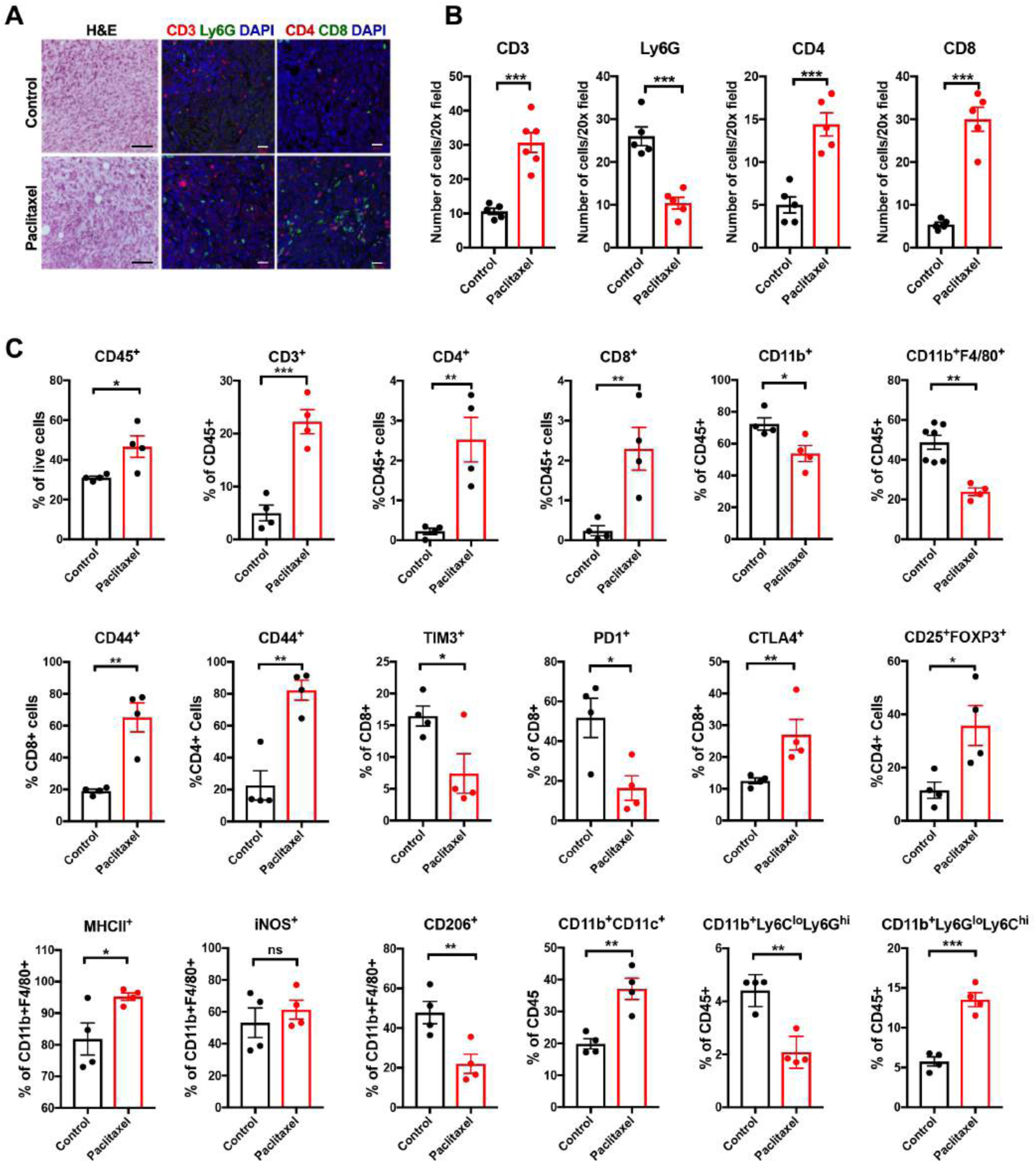
Paclitaxel Alters the TME and Reduces Tumor Burden in *Tp53*^*-/-*^;*Pten*^*-/-*^;*Nf1*^*-/-*^ FTE-derived HGSOC, related to Figure 7. **(A)** H&E and IF staining for the indicated immune markers in ovarian sections from indicated groups; black scale bars: 50 µm, white scale bars: 20 µm. (B) Quantification of the indicated immune cells in (A). Each point represents average cell number per 20X field from 5 independent sections of each mouse. Error bars indicate SEM; **P<0.01, ***P<0.001, 2-way ANOVA. (C) Flow cytometric analysis for the indicated immune subsets in *Tp53*^*-/-*^;*Pten*^*-/-*^;*Nf1*^*-/-*^ tumors with or without Paclitaxel treatment. Data indicate means ± SEM, ns, not significant, *P<0.05, **P<0.01, ***p < 0.001, unpaired t test.

**Table S1. Current collection of HGSOC organoid models and their tumorigenic potential.**

